# Coralysis enables sensitive identification of imbalanced cell types and states in single-cell data via multi-level integration

**DOI:** 10.1101/2025.02.07.637023

**Authors:** António G.G. Sousa, Johannes Smolander, Sini Junttila, Laura L. Elo

## Abstract

Current state-of-the-art integration methods for single-cell transcriptomics often struggle with imbalanced cell types across heterogeneous datasets, particularly when the datasets include similar but unshared cell types. Here, we introduce Coralysis, an R package featuring a multi-level integration algorithm to overcome these challenges. Coralysis enables sensitive integration, reference-mapping, and cell state identification across single-cell datasets, demonstrating consistent performance across diverse single-cell RNA-seq integration tasks and outperforming state-of-the-art methods when similar cell types are unevenly distributed across batches or completely absent from some datasets. Beyond single-cell transcriptomics, Coralysis enables the integration of rare cell populations from single-cell proteomic assays, such as basophils (0.5%) from whole blood. It also accurately predicts cell type identities across various query-reference scenarios. For instance, it successfully reclassifies CD16+ monocytes and natural killer cells that were previously misclassified as CD14+ monocytes and cytotoxic T cells in peripheral blood mononuclear cells. Finally, a key feature of Coralysis is its ability to provide probability scores that enable identifying both transient and steady cell states along with their differential expression programs. Overall, Coralysis facilitates the study of subtle biological variation and its dynamics by improving the integration of imbalanced cell types and states, enabling a more faithful representation of the cellular landscape in complex single-cell experiments.

## 1 Introduction

Single-cell transcriptomes from multiple subjects across different biological conditions can provide valuable insights into varying biological phenomena, such as cell trajectories^1^ in embryonic development or differential gene expression^2^ between healthy and diseased individuals. In analysing such datasets, a typical goal is to project and cluster the multi-subject single-cell transcriptomes into a low-dimensional space, ensuring that cells of the same type cluster together. However, technical noise introduced due to handling sets of samples distinctively, i.e., in batches, can distort the biological signal, leading to an unwanted outcome, where cells are clustered based on their batch identity rather than cell type. Correcting for such technical batch effect biases by integrating and “aligning” cells across batches is crucial to preserve the biological signal and accurately clustering the same cell types together.

A few benchmarks have been conducted so far to evaluate horizontal integration methods for single-cell RNA sequencing (scRNA-seq)^3–5^. They differ on the type of data used, the complexity of the integration tasks tested, and the performance metrics calculated. Tran *et al.*^3^ compared fourteen integration methods across five different integration tasks varying in complexity using four assessment metrics. They recommended Harmony^6^, LIGER^7^, and Seurat v3^8^ methods based on their overall performance. On the other hand, Richards *et al.*^4^ focused on assessing batch correction specifically for cancer scRNA-seq data integration using the LISI (Local Inverse Simpson’s Index) metric. Among the five tools tested, STACAS^9^ and fastMNN^10^ were identified as the most adequate methods regarding their ability to batch-correct the data but still retain tumour-specific biological variability. A more recent benchmark by Luecken *et al.*^5^ used the most comprehensive set of integration performance metrics to date to compare sixteen integration methods across seven scRNA-seq datasets with/without scaling and with/without feature selection. The overall three top-performing methods were scANVI^11^, Scanorama^12^ and scVI^13^. The lack of agreement between these independent benchmarks reflects that there is no single method fitting all the integration tasks. Instead, the choice of the most suitable method depends greatly on the aim of the study, the type of data, and the degree of noise introduced by the batch.

Among these methods, scANVI and scVI tools are deep neural network methods that are very effective in correcting complex batch effects,^5^ though they are less interpretable. In addition, scANVI requires cell label annotations, which are quite often absent, whereas scVI performs better with larger numbers of cells and more complex batch correction tasks^5^. In contrast, Scanorama, fastMNN, Seurat v3, and STACAS integrate the data through mutual nearest neighbours searches in a joint low-dimensional embedding, which provides the ability to remove technical noise at lower computational cost^5,14^. However, the use of a joint low-dimensional representation to integrate the batches may cause cell type “misalignment” in case there is a lack of cell type conservation across batches^3,14^. LIGER uses integrative non-negative matrix factorization to identify shared factors across batches,^7^ whereas Harmony is a linear integration method that applies iteratively *K* -means clustering in principal component space^6^. Both tend to prioritise batch correction over biological conservation,^5^ making them less suitable to integrate datasets with nuanced biological cell states.

Here, we introduce Coralysis, an R package featuring a multi-level integration algorithm for sensitive integration, reference-mapping, and cell state identification in single-cell data. Coralysis performs consistently well across diverse scRNA-seq integration tasks and query-reference scenarios. It outperforms state-of-the-art integration methods in tasks where batches do not share similar cell types, enables integration of rare cell populations beyond transcriptomics, including single-cell proteomic assays, as well as accurate reference-mapping of previously misclassified cells. Finally, a key feature of Coralysis is that it provides probability scores to identify transient and steady cell states along with their differential expression programs, providing a comprehensive approach for single-cell data integration.

## 2 Methods

### 2.1 Iterative Clustering Projection algorithm: adaptations

A multi-level integration method was implemented in Coralysis by adapting the Iterative Clustering Projection (ICP) algorithm, available in ILoReg,^15^ to work in a top-down fashion. In addition, clustering accuracy was improved and biases towards more abundant cell types and batches were effectively corrected. Coralysis is distributed as an R package and it is available at https://github.com/elolab/Coralysis. The ICP algorithm is described briefly in the next paragraph as it is at the core of our method. ICP relies mainly on two consecutive steps performed at every epoch of each of the *L* ICP runs (*L* = 200 by default): (1) logistic regression with *L*1 regularization to predict cluster probabilities; and (2) *Adjusted Rand Index* (aka *ARI*) to compare the similarity between cluster assignments across epochs. Step 1 aims to predict clusters using the ability of *L*1-regularized logistic regression to learn from the data important features and translate that into a probability matrix, *N* × *K*, with the likelihood of every *N* cell belonging to each one of the *K* clusters (*K* = 15 by default). In order to predict cluster probabilities, step 1 starts by using a balanced random initial clustering *S* where one of the *K* clusters is assigned to each *N* cell with equal probability. The training data set is created by downsampling each cell cluster to an equal number of cells. The down-sampling depends on parameter *d* (0.3 by default), meaning that 30% of the cells from each cluster are selected. An *L*1-regularized logistic regression classifier is trained on the trained data and used to classify the whole data, resulting in a new projected clustering *S^′^*. Then, step 2 compares the ARI score between the clusterings *S* and *S^′^*. If *ARI*(*S^′^, S*) increases from its previous value (initialized to zero in the first epoch), the new clustering *S^′^* is used to perform the next epoch, i.e., step 1, otherwise *S* is used to repeat step 1 until the maximum number of reiterations *r* is reached (*r* = 5 by default). Alternatively, the ARI score can increase consecutively until the maximum number of iterations *max.iter* is reached (*max.iter* = 200). Step 2 works as a measure to direct predictions towards more stable clusters and to improve the ability of the regularized regression to learn from the data. Once the best ARI value is reached for a given ICP run, i.e., the best clustering result, the algorithm jumps to the next ICP round until *L* = 200.

The adaptations introduced to the ICP algorithm will be described in the sections below.

#### 2.1.1 Training set

In order to overcome the need for using the same data for training and testing, as it was originally implemented, Coralysis builds a separate training set from the data. The process involves four steps: (1) selecting highly variable genes (HVGs) (by default 2,000) with the scran R package^16^; (2) performing a principal component analysis (PCA) using the HVGs (default PCs=30) with the irlba package^17^; (3) using the selected PCs to cluster cells into a large number of clusters (by default 500) with *k* -means++ from the package flexclust^18^; and (4) averaging gene expression within the clusters found at step (3). This results in a training set that is more balanced, less sparse and distinct from the testing data avoiding overfitting and improving the overall accuracy of classification, particularly, due to the reduced bias towards abundant cell types and batches. In addition, the ICP algorithm converges faster and uses less memory due to reduced dimensionality. In the presence of a known batch, this process is applied batch wise and the resulting datasets are concatenated into a joint training set.

#### 2.1.2 Number of clusters at start: 2

A cluster seed approach was implemented to assign every cell to one of the two clusters required at the start of the divisive ICP algorithm. The approach consists of assigning cells, in a batch wise manner, to one of the two clusters based on their distribution along the first principal component. A PCA is performed with the irlba package on the standardized data (Z-score) comprising all the cells, regardless of the batch, and all genes with a non-zero standard deviation. Then, the cells with a batch wise PC1 score lower than or equal to the median batch wise PC1 score are assigned to cluster number one and the other cells to cluster number two. The same cluster seed is given across the multiple ICP runs. This approach reduces the algorithm runtime, as it takes less epochs to converge, and it favours integration due to the more concordant clusterings obtained across multiple ICP runs. The aim of the cluster seed approach is to provide a rough, meaningful clustering result that is not robust, allowing the ICP algorithm to iterate over the clusters and converge. If a training set is built from the original data, the cluster seed process is performed over clusters of cells rather than cells.

#### 2.1.3 Cells for training: batch *k* -NN

At every epoch, except the first, the cell with the highest probability for each batch within each cluster is identified based on the current clustering and the associated probabilities. The respective *k* -nearest neighbors (*k* -NN) are then selected for training the *L1*-regularized logistic regression model with the RANN package^19^. The number of neighbours *k* can be specified as fixed or proportional to the cluster size. By default, *k* is set to 30% of the cells (or cell clusters) comprising the cluster divided by the number of batches. This selection procedure aims to find the most similar cells across batches by using probability as a proxy for similarity and to achieve a more balanced set of cells for training. Additionally, it contributes to the faster convergence of the ICP algorithm by relying on the neighbours of the cells with the highest probabilities. If a training set is used, the selection procedure of cells for training is performed over clusters of cells rather than cells.

#### 2.1.4 Divisive clustering: 2 → 4

The ICP algorithm^15^ was adapted to perform clustering in a top-down fashion by sequentially increasing the number of clusters in powers of two, i.e., 2*^k^*, where *k* ∈ Z^+^, until it reaches the target number of clusters (by default 16) - also in the power of two. The divisive clustering process begins with two clusters assigned by the cluster seed procedure described in section 2.1.2. ICP iterates over these two clusters until convergence. This process results in the assignment of a cluster label (1 or 2) and a probability score to every cell. The probability score indicates the likelihood of a cell belonging to one cluster and not the other. Next, each cluster is divided into two based on their cluster probability distribution (median as default cutoff) in a batch wise manner to avoid bias. Then ICP iterates over the four clusters until convergence. This incremental clustering process is repeated until the target number of clusters is reached (by default 16: 2 → 4 → 8 → 16). If a training set is used, the divisive clustering process is applied over clusters of cells rather than cells.

### 2.2 Coralysis reference-mapping method

The Coralysis reference-mapping method relies solely on the ICP algorithm. This means that an annotated reference trained with the divisive or the original ICP algorithm can be used to classify and project new unannotated query single-cell datasets. First, an annotated reference, i.e., with known cell types, is trained with ICP and the resulting cluster probabilities are used to perform PCA with the *stats* package^20^. Second, the cluster probabilities and PCA scores are predicted for the query or unannotated dataset(s). This procedure is dataset-independent, as cluster probabilities and PCA scores are predicted cell wise. Third, a *k* -NN classifier, from the package *class*^21^, is trained with the reference PCA and labels to transfer. Finally, the built *k* -NN model is used to classify the query cells using *k* neighbours (by default 10) and the winning class is decided by majority voting. The proportion of *k* nearest neighbours for the winning class is used as an indication of confidence. Additionally, the query PCA data can be projected onto the reference Uniform Manifold Approximation and Projection (UMAP).

### 2.3 Benchmarking integration: scib-pipeline

Coralysis integration method was benchmarked against six unsupervised integration methods (scVI, Scanorama, Seurat v4 CCA/RPCA, Harmony, fastMNN) across six datasets (pancreas, lung, human immune, human/mouse immune, two simulations) using 14 assessment metrics within the publicly available scib-pipeline (https://github.com/theislab/scib-pipeline)^5^. The mouse brain dataset was not included in the benchmark because it was not available on figshare with the other datasets due to size limitations. A detailed description of the datasets can be found in Luecken *et al.*^5^. The version of R and Seurat was updated to 4.1.3 and 4.3, respectively. The forked version of the *theislab/scib* − *pipeline* github repository with the adaptations required to reproduce the benchmark is available at: https://github.com/elolab/scib-pipeline. The benchmark was run with Snakemake (v.7.25.2) in a cluster environment with Slurm (v.23.02.6) with 8 threads and 354 GB of RAM.

### 2.4 Accuracy of reference-mapping across query-reference scenarios

The accuracy of Coralysis reference-mapping method was assessed across four different query-reference scenarios: 1) imbalanced cell types; 2) unshared cell types; 3) unrepresented batch; and 4) a varied strength of the batch effect. Performance was measured in terms of accuracy as follows:

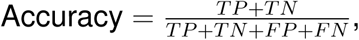

with *TP* = True Positives, *TN* = True Negatives, *FP* = False Positives, *FN* = False Negatives. These analyses were performed with the R programming language in a containerized docker image publicly available at Docker Hub (elolabfi/sctoolkit) running R (v.4.2.1)^20^ (https://www.r-project.org/) and RStudio server (2022.07.2 Build 576).

#### 2.4.1 Imbalanced cell types

Two scRNA-seq datasets (batch 1 and 2) of peripheral blood mononuclear cells (PBMCs) were used to assess the accuracy of reference-mapping for a scenario when the reference has imbalanced cell types relative to the query. The batch 1 dataset was used as reference and the batch 2 as query. Each dataset was comprised of six balanced cell types (B cell, CD4 T cell, CD8 T cell, Monocyte CD14, Monocyte FCGR3A, NK cell), i.e., each cell type had the same number of cells (n=200). In total, each dataset had 1,200 cells and 33,694 genes. The analyzed PBMC datasets are from Maan *et al.*^22^ and they can be downloaded at figshare: https://doi.org/10.6084/m9.figshare.24625302.v1. Both datasets were normalised by applying the natural-log transformation to the gene expression counts, divided by the total counts per cell multiplied by a scaling factor (10,000). The reference-mapping was performed by randomly downsampling one cell type in the reference to 10%; training the reference with ICP (2,000 HVGs); and projecting and classifying the query against the reference. This procedure was repeated independently for every cell type in the reference.

#### 2.4.2 Unshared cell types

The two PBMC datasets used for the imbalanced cell type query-reference scenario were used for assessing the performance of the reference-mapping when the reference has unshared cell types relative to the query. The reference-mapping analysis followed the same procedure described in section 2.4.1 but replacing the downsampling by ablation, i.e., completely removing one cell type at a time from the reference.

#### 2.4.3 Unrepresented batch

A set of pancreatic scRNA-seq datasets comprising eight samples (celseq, celseq2, smartseq2, fluidigmc1, indrop1, indrop2, indrop3, indrop4) sequenced across five library preparation technologies (SMARTSeq2, Fluidigm C1, CelSeq, CelSeq2, inDrops) was used to assess the performance of reference-mapping with and without shared batch effects. Every sample corresponded to a pancreatic scRNA-seq dataset sequenced using a different library preparation technology with the exception of the samples indrop1, indrop2, indrop3, indrop4 which corresponded to four biological replicates coming from the same technology inDrops. The datasets were normalised as described in section 2.4.1 and genes not expressed were removed. In total, the datasets comprised 14,890 cells and 29,340 expressed genes. The number of cells per sample varied from 638 in fluidigmc1 to 3,605 in indrop3 (celseq=1,004; indrop4=1,303; indrop2=1,724; indrop1=1,937; celseq2=2,285; smartseq2=2,394). A reference was built by integrating six samples - celseq, celseq2, fluidigmc1, indrop1, indrop3, indrop4 - using Coralysis with 2,000 highly variable genes. The samples indrop2 and smart-seq2 were chosen as query datasets representing a scenario of a shared and unshared batch effect between the reference-query, respectively. Then the query datasets were classified and projected onto the integrated reference with Coralysis. The pancreatic datasets used herein were obtained from the SeuratData package (v.0.2.1).

#### 2.4.4 A varied strength of the batch effect

Two human PBMC scRNA-seq samples representing resting and interferon-stimulated cells were used for assessing the performance of reference-mapping when the gene expression of different cell types within a sample respond distinctly to a batch effect or a biological condition, i.e., the transcriptome changes more for some cell types than others. The resting sample was chosen as reference (6,548 cells) and the interferon-stimulated sample as query (7,451). The datasets were normalised as described in section 2.4.1 and genes not expressed were removed. In total, the samples comprised 14,044 genes and 13,999 cells. The reference was trained using Coralysis with 2,000 highly variable genes and the query classified and projected onto the reference. The PBMC data used for this task was obtained from the SeuratData package (v.0.2.1).

### 2.5 Integration of PBMCs from two 10X 3’ assays

Two 10X scRNA-seq 3’ assays - V1 and V2 - from human peripheral blood mononuclear cells (PBMCs) were integrated with Coralysis to demonstrate multi-level integration in practice. Both assays were downloaded from the 10X website: V1 (https://support.10xgenomics.com/single-cell-gene-expression/datasets/1.1.0/pbmc6k) and V2 (https://support.10xgenomics.com/single-cell-gene-expression/datasets/2.1.0/pbmc8k). Cells were annotated using the annotations provided by Korsunsky *et al.* - Source Data Figure 4 file^6^. Cells without annotation were discarded (143 cells). Duplicated gene symbols were renamed by appending the corresponding Ensembl IDs. Genes not expressed were removed. In total 20,016 genes and 13,016 cells remained - 4,770 cells in V1 and 8,276 cells in V2. Normalisation was done similarly as in section 2.4.1. Highly variable genes (HVG=2,000) were selected with the package scran. The two assays were integrated using Coralysis by running the function *RunParallelDivisiveICP*, using 2,000 HVGs and default parameters, except for *allow.free.k* = *FALSE*, *C* = 1, *train.k.nn.prop* = 0.45, and *ari.cutoff* = 0.1, which were specified to ensure that Coralysis returned exactly 16 clusters. By default, *allow.free.k* = *TRUE*, allowing clusters without assigned cells to be dropped among the target number of clusters (default 16). Setting *allow.free.k* = *FALSE* overrides this behavior to enforce the return of all 16 clusters with the support of more relaxed parameters: *C* = 1, *train.k.nn.prop* = 0.45, and *ari.cutoff* = 0.1. The probability tables for each final divisive round at *K* =2, *K* =4, *K* =8, and *K* =16 were independently concatenated across the 50 ICP runs to compute a PCA and *t* -SNE with Coralysis. Probabilities were centered and scaled before PCA. The divisive ICP clustering run number two (*L*=2) corresponding to the cluster probability table (at *K16*) with the highest standard deviation was selected to exemplify the evolution of batch label mixing and cell type separation across one divisive ICP run (the algorithm was run 50 times). The top ten positive coefficients for every cluster across the four divisive ICP rounds for the run number 2 were also retrieved. The *t* -SNE and cluster tree plots were made with ggplot2 (v.3.5.1)^23^, pie charts with scatterpie (v.0.2.3)^24^ and heatmaps with ComplexHeatmap (v.2.14.0)^25^.

### 2.6 Integrating similar unshared cell type pairs

Two PBMC scRNA-seq samples, one representing resting PBMCs (CTRL) and other PBCMs stimulated with interferon (STIM), were used to highlight the ability of Coralysis to integrate similar unshared cell type pairs across batches. The *ifnb* dataset (v.3.1.0) published by Kang *et al.*^26^ was obtained from the SeuratData R package (v.0.2.1). Cells were normalized as described in section 2.4.1 and genes not expressed were removed. Two pairs of similar cell types that responded differently to interferon stimulation were selected: CD14–CD16 monocytes and CD4 naive–memory T cells. Monocytes responded more strongly to the stimulation than T cells, making them easier to integrate in this scenario. For each selected cell type pair, one cell type was retained in one sample and removed from the other to simulate a situation where batches do not share similar cell types. Specifically, CD16 monocytes (507 cells) and CD4 memory T cells (859) were retained in the CTRL sample but removed from STIM, while CD14 mono-cytes (2,147) and CD4 naive T cells (1,526) were retained in STIM but removed from CTRL. In total 14,044 genes and 9,366 cells remained (3,355 cells in CTRL and 6,011 in STIM). Finally, the integration task was run through the scib-pipeline as described in section 2.3. The *stim* and *seurat annotations* cell metadata columns were used as batch and cell type labels, respectively.

### 2.7 Integration of single-cell proteomics assays

Two types of single-cell proteomics assays were integrated with Coralysis: antibody derived tags (ADT) and cytometry by time of flight (CyTOF). The datasets were obtained from https://github.com/single-cell-proteomic/SCPRO-HI^27^. The cell type annotations used were the same as given in Koca & Sevilgen^27^. The ADT data consisted of human PBMCs from eight HIV-infected donors (P1-8) following their longitudinal response across vaccinated and unvaccinated individuals (published by Hao *et al.*^28^). Batch 1 comprised donors P1-4, and batch 2 donors P5-8. In total, the ADT dataset included 228 proteins and 161,764 cells (67,090 and 94,674 cells in batch 1 and 2, respectively). Cells were transformed using centered log-ratio (CLR) normalization prior to integration, utilizing the *NormalizeData* function from Seurat. The batch and cell type labels used were *Batch* and *cluster s*, respectively. The *celltype.l*2 labels originally given in Hao *et al.*^28^ were also highlighted.

The CyTOF data consisted of human whole blood datasets from Rahil *et al.*^29^ and Bjornson-Hooper *et al.*^30^. The dataset from Rahil *et al.*^29^ included whole blood from 35 donors infected with H1N1 influenza across 11 time points, while the dataset from Bjornson-Hooper *et al.*^30^ comprised blood samples from healthy donors exposed *in vitro* to 15 different stimuli. Only the data corresponding to the IFN*γ* stimulus was used for the latter. The dataset integrated included the expression of 39 proteins across 216,322 cells (102,147 cells from *h*1*n*1 dataset, and 114,175 cells from *ifng* dataset). The two datasets were considered as batches and the *cluster s* cell metadata column as the cell type label. The dataset was scaled (Z-score) by proteins before integration. The first seven principal components were used for building the training set instead of the default 30 due to the low number of features (*n*=39).

In both integration tasks, all features were used for integration and the resulting probabilities were centered and scaled before PCA with Coralysis. In addition, the fast implemented version of UMAP available in the uwot (v.0.1.14)^31^ R package was used to compute UMAP. The python package scib-metrics^5^ was used to compute batch-correction and bio-conservation metrics without the calculation of isolated labels as the batches shared all cell types.

### 2.8 Mapping PBMCs from different sequencing technologies

The *pbmcsca* scRNA-seq dataset (v.3.0.0) from SeuratData was used to quantify the mapping accuracy performance of Coralysis. The dataset published by Ding *et al.*^32^ comprised nine PBMC samples prepared with different single-cell library preparation technologies: 10x Chromium (v2) A, 10x Chromium (v2), 10x Chromium (v2) B, 10x Chromium (v3), CEL-Seq2, Drop-seq, inDrops, Seq-Well, Smart-seq2. The sample corresponding to PBMCs obtained through 10x Chromium (v2) A (3,222 cells) was selected as a reference and the remaining as query datasets (27,753 cells). Cells without annotation, i.e., with label *Unassigned* in *CellType* cell metadata column, were removed. The data were normalised as described in section 2.4.1 and genes not expressed were discarded. The top 2,000 most highly variable genes were selected with the scran function *modelGeneV ar* for the reference sample (10x Chromium (v2) A). The reference sample was trained with Coralysis function *RunParallelDivisiveICP*, with default parameters, with the exception of *divisive.method* = “*cluster*”, as the reference sample did not have cells originating from different batches, i.e., *batch.label* = *NULL*. The PCA and UMAP were computed with Coralysis with the argument *return.model* = *TRUE*. Probabilities were scaled prior to PCA as described in section 2.5. Reference-mapping was computed with Coralysis function *ReferenceMapping* with default parameters and cells were projected onto the reference UMAP (*project.umap* = *TRUE*). Visualizations were obtained with Coralysis, ggplot2 (v.3.5.1), and scater (v.1.26.1)^33^.

### 2.9 Inference of cell states with cell cluster probabilities

CD34+ hematopoietic stem cells from Persad *et al.*^34^ were used to show how Coralysis cell cluster probability can be biologically meaningful. The scRNA-seq dataset consisted of 6,881 cells downloaded from Zenodo: https://zenodo.org/records/6383269/files/cd34_multiome_rna.h5ad. The top 2,000 highly variable genes were selected with the scran function *getTopHV Gs* before running Coralysis function *RunParallelDivisiveICP* with *divisive.method* = “*cluster*”. UMAP was calculated with the uwot package and parameters were adjusted to provide more connectivity: *n neighbors* = 100, *min dist* = 0.5. The *palantir pseudotime*^35^ variable was used to compute the Pearson correlation with the mean cell cluster probability obtained with Coralysis across 50 ICP runs. The cell type labels used corresponded to those available in the *celltype* variable as used in the original study.

Human embryoid body cells (31,029 cells) from Moon *et al.*^36^ were integrated by embryonic stage (0-1, 2-3, 4-5, 6-7, 8-9) with Coralysis using 6,000 HVGs. UMAP was computed as described above for the CD34+ cells dataset. The Coralysis cell cluster probabilities were divided into 20 evenly sized bins for each cell type independently. Gene expression and cell cluster probabilities were then averaged within each bin for each cell type to calculate the Pearson correlation between them.

Visualisations were obtained with ggplot2 (v.3.5.1) and ComplexHeatmap (v.2.14.0).

### 2.10 Data availability

All datasets analysed in this study have been previously published and made publicly available by the original authors. Links to each dataset are provided in the methods section alongside their descriptions.

### 2.11 Code availability

The multi-level integration and reference-mapping methods have been implemented into the R package Coralysis publicly available on GitHub: https://github.com/elolab/Coralysis. The forked version of the *theislab/scib* − *pipeline* github repository with the adaptations required to reproduce the benchmark described herein is available at: https://github.com/elolab/scib-pipeline. Finally, all the R and python scripts required to reproduce the remaining analyses and figures are available at: https://github.com/elolab/Coralysis-reproducibility.

## 3 Results

### 3.1 Coralysis identifies cell types through multi-level integration

Coralysis identifies shared cell subpopulations (cell types or states) across a given set of heterogeneous single-cell datasets through four main steps (see Extended Data Fig. 1A). The first step clusters the cells batch-wise in a low dimensional space (PCA) using Kmeans++^18^ with a large number of clusters (default 500). These clusters are used to average the feature expression in order to build a training set. The second step uses the built training set to identify shared cell clusters across datasets through divisive ICP modelling. In the third step, the cluster probability is predicted for every cell against every ICP model. Finally, the fourth step concatenates the cluster probability matrices and performs a PCA to determine an integrated embedding, which can be used for downstream clustering or non-linear dimensionality reduction techniques.

The original ICP algorithm was not designed for clustering heterogeneous scRNA-seq datasets, and consequently, it tends to emphasise technical noise or batch effects, despite the implemented ensemble clustering approach. Therefore, we adapted here the ICP algorithm to perform divisive clustering rounds in order to promote the batch label mixture and cell type separation (Extended Data Fig. 1B-E). Briefly, we begin with two clusters determined by the cell distribution across the first principal component, in a batch-wise manner (Extended Data Fig. 1B). For every epoch, i.e., a single model iteration through the training set, the ICP algorithm selects for training the clusters those cells that represent the batch-wise *k* nearest neighbours with the highest probability (Extended Data Fig. 1C). Once the algorithm converges to a stable clustering, the next clustering round starts by splitting every cluster obtained into two based on the batchwise cluster probability distribution (Extended Data Fig. 1D). The process (Extended Data Fig. 1C-D) is repeated until the algorithm reaches the last clustering round (by default clustering round four, corresponding to 16 target clusters; Extended Data Fig. 1E).

To demonstrate how the Coralysis multi-level integration works in practice, we used two 10X scRNA-seq datasets of peripheral blood mononuclear cells (PBMCs) obtained with two different 3’ assays (V1 and V2). The dataset V2 comprised ≈1.7 times more cells than V1 (8,276 versus 4,770 cells), making this an unbalanced integration task, which are typically more challenging (Fig. 1A). The divisive ICP clustering run number two (*L*=2) corresponding to the cluster probability table (at *K16*) with the highest standard deviation was selected to exemplify the evolution of batch label mixing and cell type separation across one divisive ICP run (the algorithm was run 50 times). As expected, the median cluster probability decreased throughout the divisive ICP clustering rounds, i.e., *K*2 → *K*4 → *K*8 → *K*16 (Fig. 1B). The batch label distribution per cluster was dominated by cells coming from the assay V2, as expected, given the fact it comprised almost twice the number of cells compared to assay V1 (Fig. 1C). The only exception was cluster 8, which consisted mainly of CD16+ monocytes at *K16* (Fig. 1D). Interestingly, CD16+ monocytes were the only cell type with more cells in dataset V1 than V2 (332 versus 225, respectively). This highlights the ability of Coralysis to handle the integration of unbalanced datasets and cell types.

**Figure 1.**
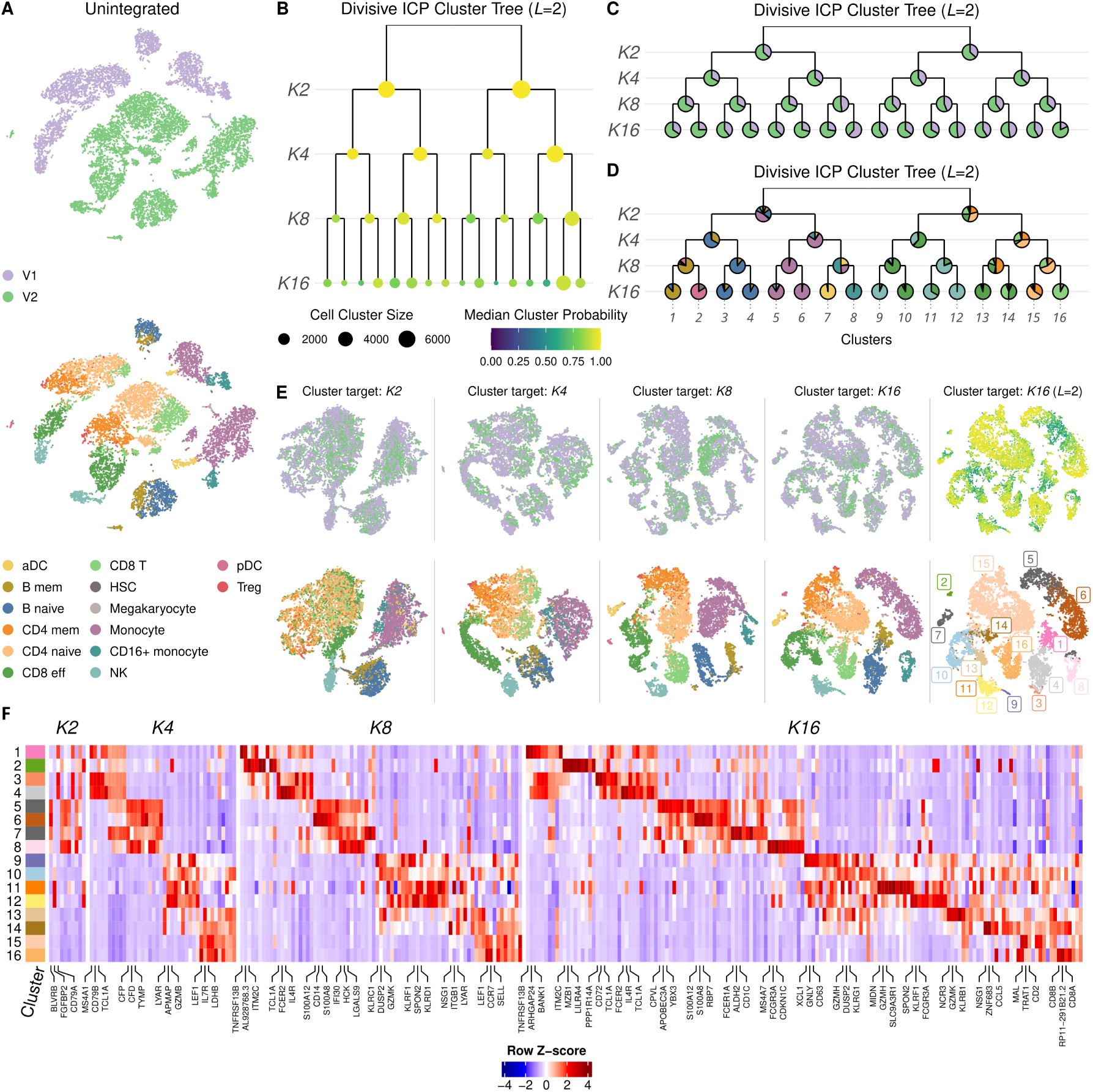
Demonstration of Coralysis multi-level integration method. **A** Unintegrated peripheral blood mononuclear cells from two 10X 3’ assays, V1 (*n*=4,770 cells) and V2 (*n*=8,276 cells), projected onto *t* -SNE comprising activated dendritic cells (aDC), B memory cells (B memory), B naive cells, CD4 memory cells (CD4 mem), CD4 naive cells, CD8 effector T cells (CD8 eff), CD8 T cells (CD8 T), hematopoietic stem cells (HSC), megakaryocytes, monocytes, CD16^+^ monocytes, natural killer cells (NK), plasmacytoid dendritic cells (pDC) and regulatory T cells (Treg). **B** Divisive ICP clustering tree for the divisive ICP run corresponding to the *K16* cluster probability table with the highest standard deviation (*L*=2). Size of the dots represents cluster size and the colour the median cluster probability. **C** Cell distribution of batches per cluster for the divisive ICP run 2. **D** Distribution of cell types per cluster for the divisive ICP run 2. **E** *t* -SNE projections demonstrating multi-level integration in practice highlighting the fade of batch effect and separation of cell types along the four divisive ICP rounds (2, 4, 8 and 16). **F** Top ten positive coefficients for every cluster across the four divisive ICP rounds for the run 2. Rows represent the 16 clusters found at divisive ICP run 2 and columns coefficients, i.e., genes. The normalized gene expression data was averaged across the clusters and standardized (Z-score) cluster wise. The top three coefficients for every cluster at every round are labeled.

The clusters at *K2* separated monocytes, dendritic cells (DCs), natural killer (NK), and B cells from T cells (Fig. 1D). This separation was also observed in the *t* -SNE projection of the concatenated cluster probability matrices at *K2* across the 50 independent ICP runs (Fig. 1E). Noticeably, the expression of the top three positive gene coefficients for the ICP model corresponded to *BLVRB*, *FGFBP2* and *CD79A*, confined to monocytes/DCs, NK, and B cells, respectively (Fig. 1F, Extended Data Fig. 2A and Supplementary Table 1).

At *K4*, the first cluster gives rise to a B cell (naive and memory) and a monocyte/DC cluster (CD14+ and CD16+ monocytes and activated DCs) (Fig. 1D and F). The second cluster is partitioned into an effector CD8/NK and a CD8/CD4 (naive and memory) T cell cluster. This division is supported by coefficients of the ICP model, including well-known cell type-specific genes for B cells (*MS4A1*, *CD79B/A*, *TCL1A*, *LINC00926*, HLA class II histocompatibility antigen genes); monocytes/DCs (*CFP*, *CFD*, *TYMP*, *AIF1*, *FCN1*, *S100A8*, *SAT1*, *LST1*, *CST3*, *CSTA*); effector CD8/NK cells (*LYAR*, *APMAP*, granzymes (GZMB/H/M), *FGFBP2*, *CMC1*, chemokines (*CCL5*, *XCL2*), *CTSW*); and CD4/8 naive/memory cells (*LEF1*, *IL7R*, *LDHB*, *NOSIP*, *LDLRAP1*, *LTB*, *NPM1*, *SPOCK2*, *IL32*, *CD3D*) (Fig. 1F, Extended Data Fig. 2B and Supplementary Table 1).

At *K8*, B memory and plasmocytoid DCs separated from B naive cells, and similarly CD14+ monocytes separated from CD16+ monocytes and activated DCs (aDCs) (Fig. 1D-F, Extended Data Fig. 2C and Supplementary Table 1). The other four clusters involved mostly CD8 effector cells, NK cells, CD4 memory cells, and CD4 naive cells, respectively. At *K16*, the clusters were further divided into separate B memory cells (*MS4A1*, *CD79A/B*) from pDCs (*IRF7*, *XBP1*, *LILRA4*) and aDCs (*FCER1A*, *CD1C*) from CD16+ monocytes (*FCGR3A*) despite the uneven number of cells comprising these cell types (Fig. 1D-F, Extended Data Fig. 2D and Supplementary Table 1). Transcriptionally similar cell types were also further separated, such as NK cells (*GNLY*, *KLRD1*, *GZMK*, *CD63*) from CD8 effector cells (*GZMH/K/M*, *KLRG1*, *CCL5*, *CST7*). These results demonstrate the sensitivity of Coralysis to integrate cell populations by selecting low-to-high resolution cell type-specific gene coefficients along the top-down clustering process.

### 3.2 Coralysis prioritises bio-conservation

We benchmarked Coralysis using the publicly available scib-pipeline^5^ in order to perform an independent and unbiased comparison. The top five unsupervised integration methods from the benchmark conducted by Luecken *et al.*^5^ were selected for the comparison (scVI^13^, Scanorama^12^, Seurat v4 RPCA^28^, Harmony^6^ and fastMNN^10^). Additionally, we included the widely used Seurat v4 CCA method^28^. The methods were benchmarked across four real datasets (pancreas, lung atlas, human immune, human/mouse immune) and two simulations. We used the same type of input (with and without highly variable gene (HVG) selection and with or without scaling) and output (batch-corrected gene expression matrix and/or embedding), datasets (*n*=6), and performance metrics (*n*=14) as the original benchmark study^5^. In total, the scib-pipeline ran 210 tasks. Eleven of these failed: Coralysis given the HVG unscaled gene expression matrix as input for the simulation 1 dataset; Seurat v4 CCA given the full scaled gene expression matrix as input for the lung atlas and human/mouse immune datasets; and all the Seurat v4 RPCA tasks for the human/mouse immune and simulation 2 datasets (Extended Data Fig. 3A-B).

The trade-off between batch correction and bio-conservation was assessed by averaging the five batch correction and the nine bio-conservation metrics for each input data and each integration method (Extended Data Fig. 3A). Coralysis performed consistently well across the different datasets, demonstrating a competitive and balanced trade-off between batch correction and biological signal preservation (Extended Data Fig. 3A-C- and Supplementary Fig. 1-6). In general, the differences observed in the performance between the datasets reflected the complexity of the integration tasks. The human pancreas data consisted of the same tissue (pancreatic islets) analysed using distinct single-cell library preparation methods, the lung atlas included transcriptionally similar but functionally different cell types from the lung airway and parenchyma (Supplementary Fig. 7A-B), while the human and cross-species immune datasets represented more complex integration challenges with multiple batches, tissues (blood and bone marrow) (Supplementary Fig. 7C), and even organisms (human and mouse) (Supplementary Fig. 7D). Among the two simulation scenarios, the simulation 2 included more batches and fewer clusters of transcriptionally similar identities than simulation 1 (Supplementary Fig. 8A-B). Thus, it is not surprising that the benchmarked methods performed generally better in simulation 1 than in simulation 2 (Extended Data Fig. 3A-B and Supplementary Fig. 5,6).

Another important aspect of Coralysis is the small variation in its performance with respect to the input provided, compared with the other methods (Extended Data Fig. 3D). Importantly, Coralysis avoided the integration of similar yet distinct cell types (Extended Data Fig. 4-5). For example, in the human immune data, Coralysis was able to separate CD14+ from CD16+ monocytes and NK from NKT cells (Extended Data Fig. 4C). Higher bio-conservation metrics of Coralysis, such as cell type ASW (average silhouette width) and isolated labels (F1 and silhouette), (Extended Data Fig. 3B) supported these observations. Cell type ASW measures the density and separation of cell types, while the isolated label metrics assess the ability to integrate cell types shared by only a few batches. This highlights the potential of Coralysis to outperform current top-performing integration methods in tasks where subtle biological variation and/or uneven distribution of cell types across batches need to be integrated.

### 3.3 Coralysis outperforms in integration of unshared similar cell types

To further investigate the hypothesis that Coralysis outperforms the top-performing integration methods in tasks involving uneven or unshared, but transcriptionally similar cell types, we envisioned three possible scenarios for unshared cell types across batches (Fig. 2A): cell type unique to one of the batches, cell types unique to both batches that are transcriptionally distinct, and cell types unique to both batches that are transcriptionally similar. Since Scanorama, Seurat v4 CCA/RPCA, fastMNN, and Harmony perform integration in the joint low-dimensional space, they may overlook subtle biological differences that are not well preserved in a reduced feature space, compared to probabilistic models like Coralysis or scVI. Accordingly, although integration tasks involving the first two scenarios are unlikely to be wrongly integrated, it is likely that transcriptionally similar cell types will incorrectly correspond to mutual nearest neighbours.

**Figure 2.**
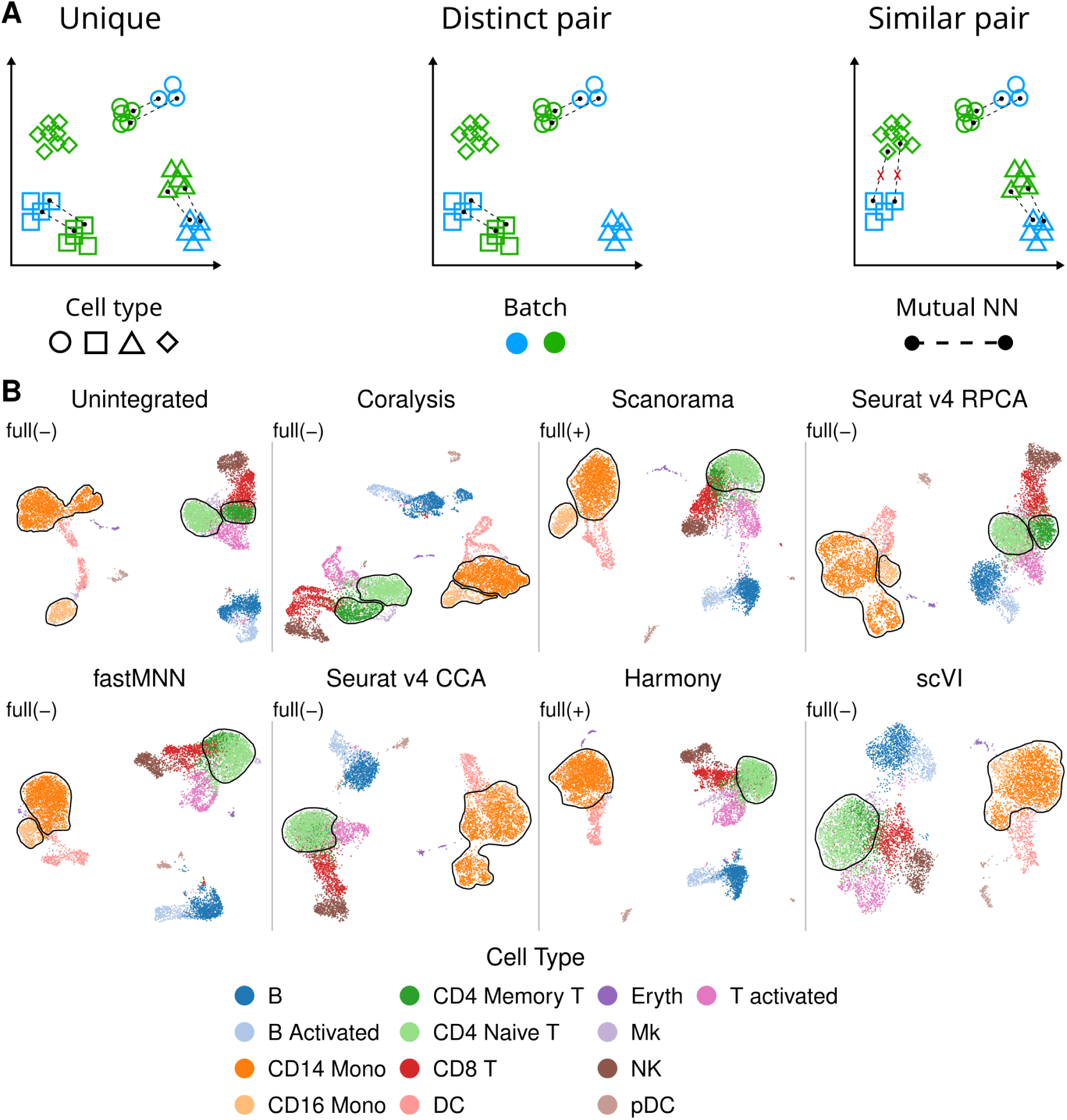
Coralysis outperforms top-performer methods for an integration task with batches with unshared similar cell type pairs. **A** Representation of mutual nearest neighbours search on the low dimensional space for integration scenarios where batches do not share all cell types: unique (one cell type is unique to one of the batches), distinct pair (one distinct pair of cell types unshared across batches, e.g., monocytes versus CD4 T cells), and similar pair (one similar pair of cell types unshared across batches, e.g., CD4 naive versus CD4 memory). **B** UMAP projections highlighting the cell-type identity before and after integration of two PBMC scRNA-seq datasets through the scib-pipeline. The two PBMC datasets consisted of one sample representing resting PBMCs (CTRL) and the other interferon-stimulated cells (STIM). CD4 naive T cells and CD14 monocytes were removed from the CTRL batch sample and CD4 memory T cells and CD16 monocytes from the STIM sample. Unshared similar cell type pairs are circumscribed by black lines. The best input-output combination was selected for every method. The label “full” represents input data with all features, and the minus and plus signs correspond to unscaled and scaled data, respectively.

We investigated this hypothesis using a dataset of two human PBMC samples: resting cells (CTRL) and interferon-stimulated cells (STIM) (Fig. 2B and Supplementary Fig. 9). Specifically, we considered two pairs of similar cell types CD4 naive and CD4 memory T cells, and CD14 and CD16 monocytes. In the CTRL sample, we retained CD16 monocytes and CD4 memory T cells but dropped out CD14 monocytes and CD4 naive T cells. In the STIM sample, we retained and dropped out vice versa. Coralysis was the only method that could clearly separate CD14 from CD16 monocytes as well as naive from memory CD4 T cells (Fig. 2B and Supplementary Fig. 9,10). Despite using a probabilistic model, scVI performed poorly, contrary to our expectations (Fig. 2B and Supplementary Fig. 9,10). This result supports our hypothesis, showing that Coralysis is a more suitable method for integrating unshared similar cell types.

### 3.4 Coralysis integrates heterogeneous single-cell proteomics datasets

Although Coralysis has been developed with the purpose of integrating and/or clustering single-cell transcriptomic datasets, the method can also be applied to other single-cell modalities. To demonstrate this, we decided to apply Coralysis integration on single-cell proteomic data from two fast-evolving technologies: antibody derived tags (ADT) and cytometry by time of flight (CyTOF). The ADT data consisted of human PBMC samples from eight HIV-infected donors (P1-8) published by Hao *et al.*^28^, collected along three time points to follow both vaccinated and unvaccinated individuals (228 proteins across 161,764 cells). The normalised data was integrated across two batches: the first comprising donors P1-4, and the second, donors P5-8 (Fig. 3A), as defined in Koca & Sevilgen^27^, based on the eight higher-level cell type labels available from the same study (Fig. 3B), Coralysis integrated all cell populations well (Fig. 3C-D). The high granularity observed independent of the batch (Fig. 3C,E) motivated us to further look into the more granular cell populations using the fine-grained cell type annotations from the original study (at level 2 of granularity), which had been obtained using CITE-seq with both ADT plus RNA-seq data modalities^28^ (Fig. 3E,F). Interestingly, several of the Louvain clusters identified with the Coralysis integrated embedding using only the ADT data matched the fine-grained cell populations identified with CITE-seq multi-modal data (Fig. 3F,G), for example, clusters 13 (MAIT), 21 (CD16 Mono), 23 (pDC), 18 plus 24 (gdT), and 30 (platelet) (Fig. 3F,G). Furthermore, despite the reduced feature space (228 proteins), Coralysis was able to preserve the differentiation trajectory, from naive to effector cell states, observed in B cells and CD4/CD8 T cells (Fig. 3F). These results were supported by the quantification of objective bio-conservation and batch-correction metrics, showing a great improvement in batch correction without compromising bio-conservation for any of the ground-truth cell type labels (Fig. 3H and Supplementary Fig. 11).

**Figure 3.**
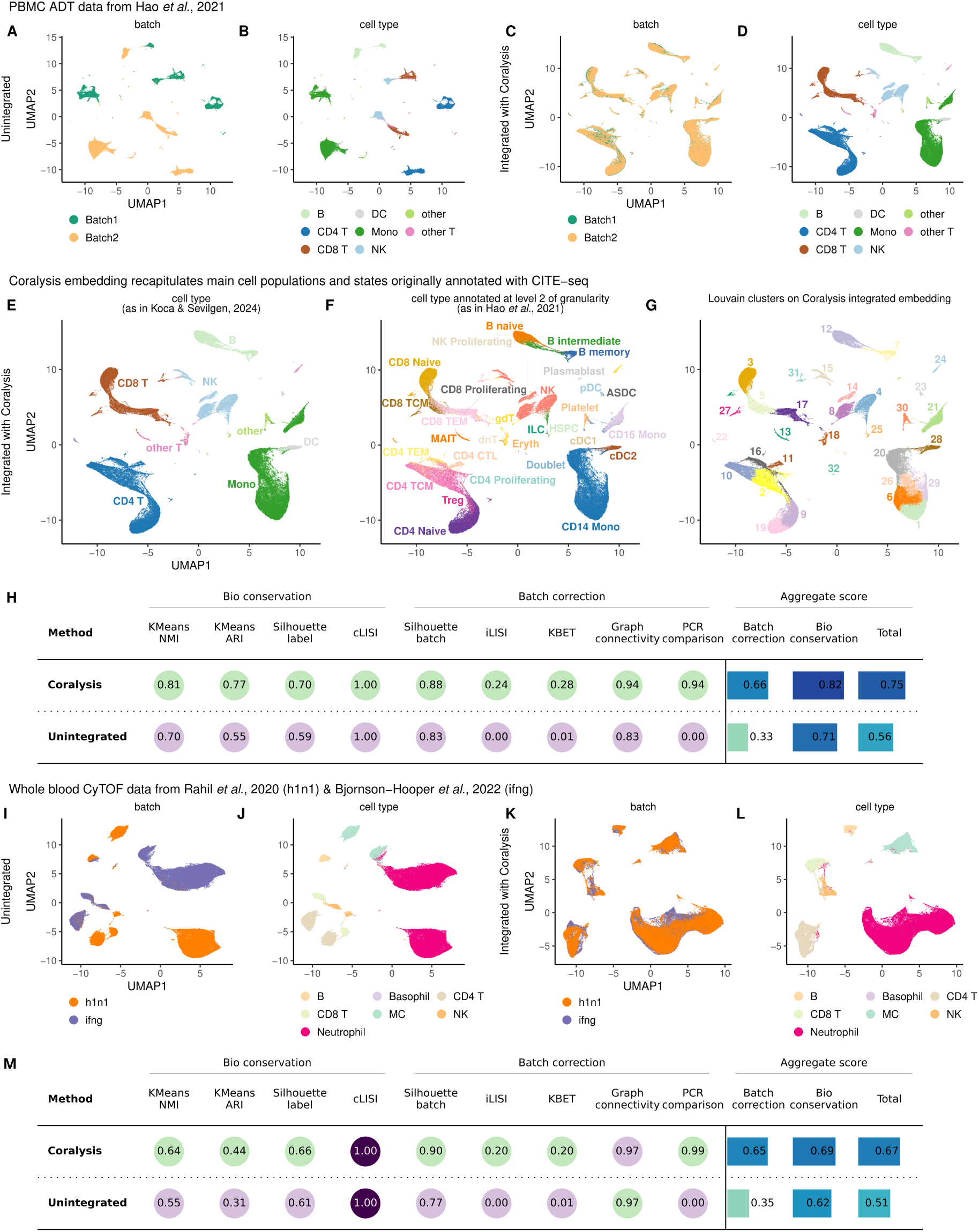
Coralysis enables the integration of heterogeneous single-cell proteomics datasets from different technologies. UMAP of unintegrated antibody derived tags (ADT) data of human PBMCs originally published by Hao *et al.*^28^ highlighting **A** the batch and **B** the cell type label identities as given in Koca & Sevilgen^27^. UMAP of the same ADT data of human PBMCs after performing integration with Coralysis highlighting **C** the batch and **D** the cell type label identities. Integrated UMAP with cells labeled as in **E** Koca & Sevilgen^27^, as in **F** Hao *et al.*^28^, at level 2 of granularity, and **G** with the Louvain clusters computed on the Coralysis integrated embedding. **H** Assessment of integration performed with Coralysis on the ADT data set provided by the scib-metrics python package using as ground-truth the cell type labels given in Koca & Sevilgen^27^. UMAP of unintegrated cytometry by time of flight (CyTOF) data of whole blood from patients infected with influenza (h1n1) or cells stimulated with interferon (ifng) originally published by Rahil *et al.*^29^ and Bjornson-Hooper *et al.*^30^, respectively, highlighting **I** the batch and **J** the cell type label identities as given in Koca & Sevilgen^27^. UMAP of the same CyTOF data of human blood cells after performing integration with Coralysis highligthing **K** the batch and **L** the cell type label identities. **M** Assessment of integration performed with Coralysis on the CyTOF dataset with the scib-metrics package.

The CyTOF datasets consisted of human whole blood samples published by Rahil *et al.*^29^ and Bjornson-Hooper *et al.*^30^, including 35 donors infected with H1N1 influenza across 11 time points (h1n1 dataset, 39 proteins across 102,147 cells), and samples from 86 healthy individuals stimulated with IFN*γ* (ifng dataset, 39 proteins across 114,175 cells) (Fig. 3I,J). As shown in the integrated UMAP projections (Fig. 3K,L), Coralysis successfully integrated all immune cell types, including the less abundant basophils (0.5% of the cells - 1,075 cells), across the highly heterogeneous CyTOF datasets. These results were further supported by the improved bio-conservation and batch-correction scores when compared to the unintegrated baseline (Fig. 3M). Over-all, these results demonstrated that Coralysis can accurately integrate single-cell proteomics data, even with a low number of features, such as CyTOF.

### 3.5 Coralysis achieves high reference-mapping accuracy under diverse query-reference scenarios

The ICP models generated by training a single-cell dataset with Coralysis can be used to predict the cluster identities of related, unannotated single-cell datasets, thereby enabling reference-mapping. To understand the strengths and limitations of Coralysis reference-mapping method, we assessed its accuracy across four query-reference scenarios: 1) imbalanced cell types, 2) unshared cell types, 3) unrepresented batches, and 4) a varied strength of the batch effect.

To assess the performance of Coralysis under the first two scenarios of imbalanced and unshared cell types, two PBMC samples from Maan *et al.*^22^ were used; the batch 1 sample of the data was used as the reference, and batch 2 as the query. Both samples consisted of six cell types (B cells, CD4 T cells, CD8 T cells, CD14 monocytes, FCGR3A monocytes, and NK cells), with each cell type containing the same number of cells (*n* = 200). For the imbalanced cell-types scenario, we iteratively down-sampled each cell type in the reference to 10% and predicted the cell type labels by projecting against the reference after training with ICP (Extended Data Fig. 6A). The centroid cross-batch distance (Euclidean) in the PCA space between the reference and query cell types was quantified to highlight their differences across the cell types (Extended Data Fig. 6B). This is particularly important since the Coralysis reference-mapping method relies on the distances between query cell types projected onto the reference PCA to classify cell types. As expected, the closest query cell-type centroid to each reference cell-type centroid corresponded to the cell type matching the reference cell-type label (Extended Data Fig. 6B). The overall accuracy ranged between 85-96% (Extended Data Fig. 6C), being lowest for CD8 T cells and highest for B cells (Extended Data Fig. 6D-E). The accuracies were consistently lower for the down-sampled cell types, particularly for CD8 T cells (Extended Data Fig. 6C and Supplementary Fig. 12A-F). Most of the CD8 T cell misclassifications were against CD4 T cells (Extended Data Fig. 6D), likely due to their transcriptional similarity, which was reflected in the smallest centroid cross-batch distance observed between the reference and query cell types (Extended Data Fig. 6C). Interestingly, the lowest Coralysis confidence scores for this classification task, corresponding to the proportion of *K* neighbors from the winning class, were associated with the CD8 T cell misclassifications (Extended Data Fig. 6D).

In the unshared cell-types scenario, we conducted a similar analysis but, instead of downsampling, each reference cell type was iteratively ablated (Extended Data Fig. 7A-B). The classification accuracy dropped compared to the previous scenario, ranging between 80-82% (Extended Data Fig. 7C), primarily due to the absence of the ablated cell type in the reference. This led to misclassifications against the closest reference cell type (Extended Data Fig. 7B and Supplementary Fig. 13A-F). Overall accuracy was lowest when B cells were ablated, and highest when CD14 monocytes were ablated (Extended Data Fig. 7D-E). In the absence of these cell types in the reference, B cells and CD14 monocytes were mostly classified against the closest reference cell-type centroids, CD4 T cells and FCGR3A monocytes (also known as CD16 monocytes), respectively (Extended Data Fig. 7B,D-E).

In the unrepresented batch scenario, we built a reference of pancreatic islets by integrating six pancreatic scRNA-seq datasets from several technologies with Coralysis (Fig. 4A-B). The queries consisted of two pancreatic scRNA-seq datasets: indrop2 (1,724 cells), which was represented in the reference, and smartseq2 (2,394 cells), which was absent from the reference (Fig. 4C-D). The prediction accuracy was very high (approximately 98%), regardless of batch representation in the reference (Fig. 4E-H). Similarly, the confidence scores were high (Fig. 4E,G).

**Figure 4.**
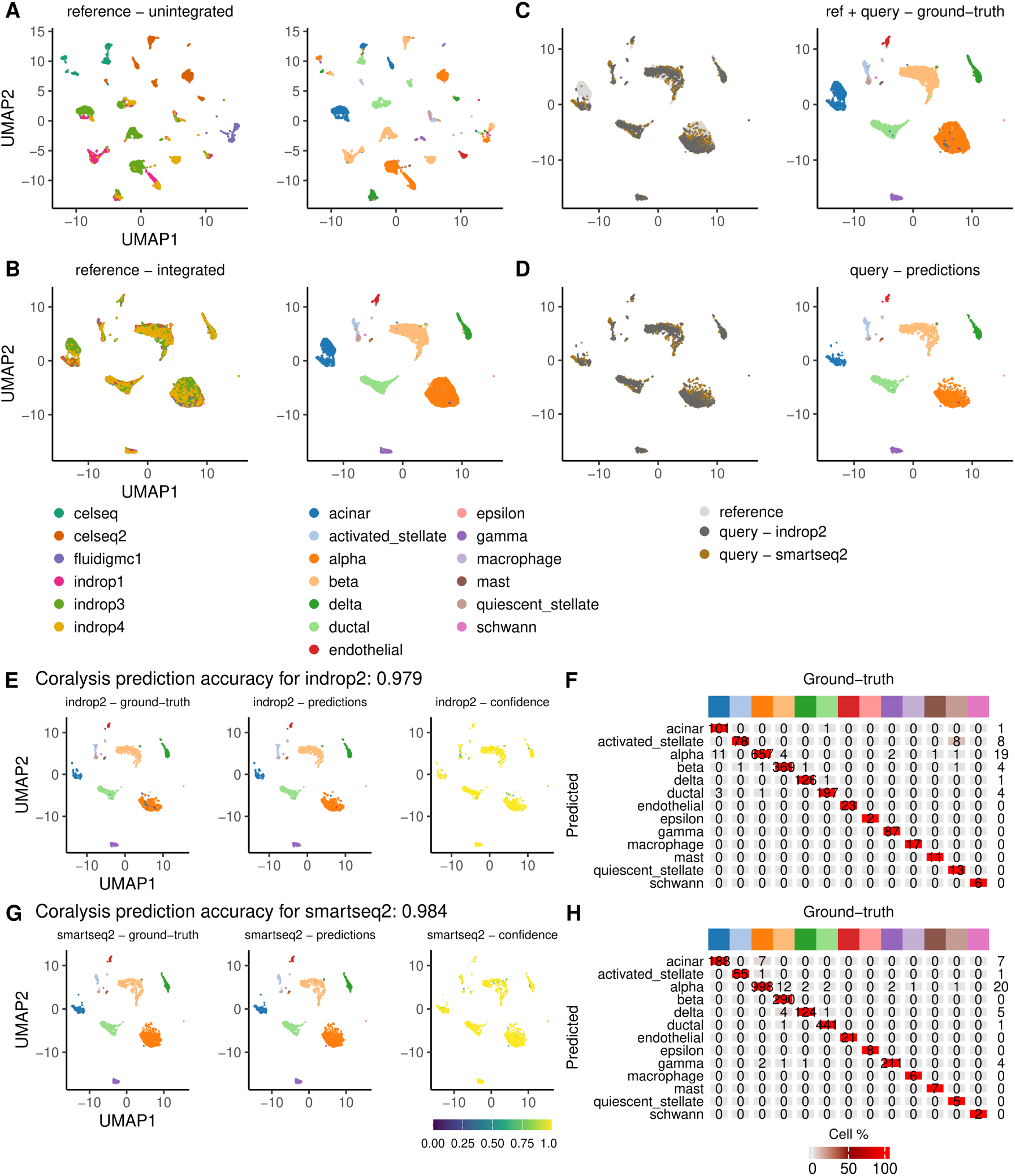
Accuracy performance of Coralysis reference-mapping method for a query-reference scenario of an unrepresented batch. **A** Unintegrated pancreatic reference UMAP highlighting the batch - sequencing libraries - and cell type labels (left-right). **B** Integrated pancreatic reference UMAP obtained with Coralysis highlighting the respective batch and cell type labels (left-right). **C** Queries projected onto the reference integrated UMAP with Coralysis reference-mapping method. Colours highlight dataset identity, i.e., reference and query (indrop2 and smartseq2) labels (left), and ground-truth cell-type labels (right). **D** Queries projected onto the reference UMAP with reference cells removed for clarity. Colours highlight query identity and predicted cell labels (left-right). UMAP highlighting ground-truth, predictions and confidence scores for **E** indrop2 and **G** smartseq2 queries projected onto the reference UMAP. Confusion matrix of predicted versus ground-truth cell labels for **F** indrop2 and **H** smartseq2 query predictions. Top colour bar represents cell type labels. The confidence scores represent the proportion of *K* neighbours from the winning class (*K* =10). The values in the confusion matrices correspond to the number of cells matching each other and the values at the end of each row, the number of total misclassifications for every predicted cell type. The heatmap colors in the matrices represent the frequency of predicted cell types in percentage.

Finally, we tested the impact of varying strengths of batch effect on the reference-mapping accuracy of Coralysis. The same resting and interferon-stimulated PBMC samples from Kang *et al.*^26^ described in section 3.3 were chosen to represent this scenario. The reference consisted of resting cells (6,548 cells) and the query of interferon-stimulated PBMCs (7,451 cells). A PCA comprising the cells from both samples was computed to highlight the transcriptomic variation between cell types (Extended Data Fig. 8A). The variation observed along Principal Component 2 (PC2) was explained by the transcriptomic response to interferon stimulation, which was stronger for monocytes and DCs than for NK, T, and B cells (Extended Data Fig. 8A). To illustrate this more clearly, the distances between the reference and query cell type centroids were depicted in a heatmap, highlighting the cell type-to-cell type distances (Extended Data Fig. 8B). Monocytes and DCs were the most dissimilar cell types, showing a clear separation from the remaining cell types, which were more similar to each other (Extended Data Fig. 8B). The correspondence between ground-truth cell labels and Coralysis predictions was high, with an overall accuracy close to 90%. The exception was a fraction of CD4 memory T cells that were wrongly predicted as CD4 naive T cells (Extended Data Fig. 8C-F). Importantly, the accuracy of classification for cell types that responded more strongly to interferon stimulation, such as monocytes and DCs (Extended Data Fig. 8A), remained high. Instead, the accuracy depended more on the similarity between the query and reference cell types (Extended Data Fig. 8B), with CD4 memory T cells being most difficult to predict due to their transcriptional similarity with CD4 naive T cells (Extended Data Fig. 8E-F). The low confidence scores in the overlapping neigh-bourhoods highlighted those cell predictions that should be more carefully inspected (Extended Data Fig. 8F). In summary, Coralysis accurately maps cells across distinct query-reference scenarios, as long as the reference faithfully represents the query.

### 3.6 Coralysis identifies rare cell populations previously missed in PBMCs

We next sought to take a closer look at the ability of Coralysis to identity fine-grained cell populations across heterogeneous scRNA-seq datasets. For this purpose, we selected nine PBMC datasets sequenced using different technologies^32^. As the reference, we selected one of the samples, which was sequenced using 10x Chromium (v2) A, whereas the remaining eight samples were used as queries (sequenced using 10x Chromium (v2), 10x Chromium (v2) B, 10x Chromium (v3), CEL-Seq2, Drop-seq, inDrops, Seq-Well, Smart-seq2). The 27,753 query cells were successfully mapped onto the 3,222 reference cells (Fig. 5A), with only minor disagreement between the ground-truth (Fig. 5B) and predicted cell labels (Fig. 5C). The mean accuracy per query dataset was 84% (Fig. 5D). Not surprisingly, all the 10x Chromium samples were above the mean as they came from the same technology as the reference. The accuracy decreased gradually from the droplet-based (Drop-seq, inDrops) to the plate-based datasets (Smart-seq2, CEL-Seq2, Seq-Well) (Fig. 5D). The distribution of confidence scores for correct and incorrect classifications was similar across datasets. Correct classifications showed a high peak close to one, whereas the incorrect classifications reached a smaller peak right after 0.5, generally followed by a uniform distribution towards one (Fig. 5E). These results support the use of confidence scores as a measure of classification reliability, particularly for scores below 0.75, where the distribution of incorrect classifications exceeds that of correct classifications.

**Figure 5.**
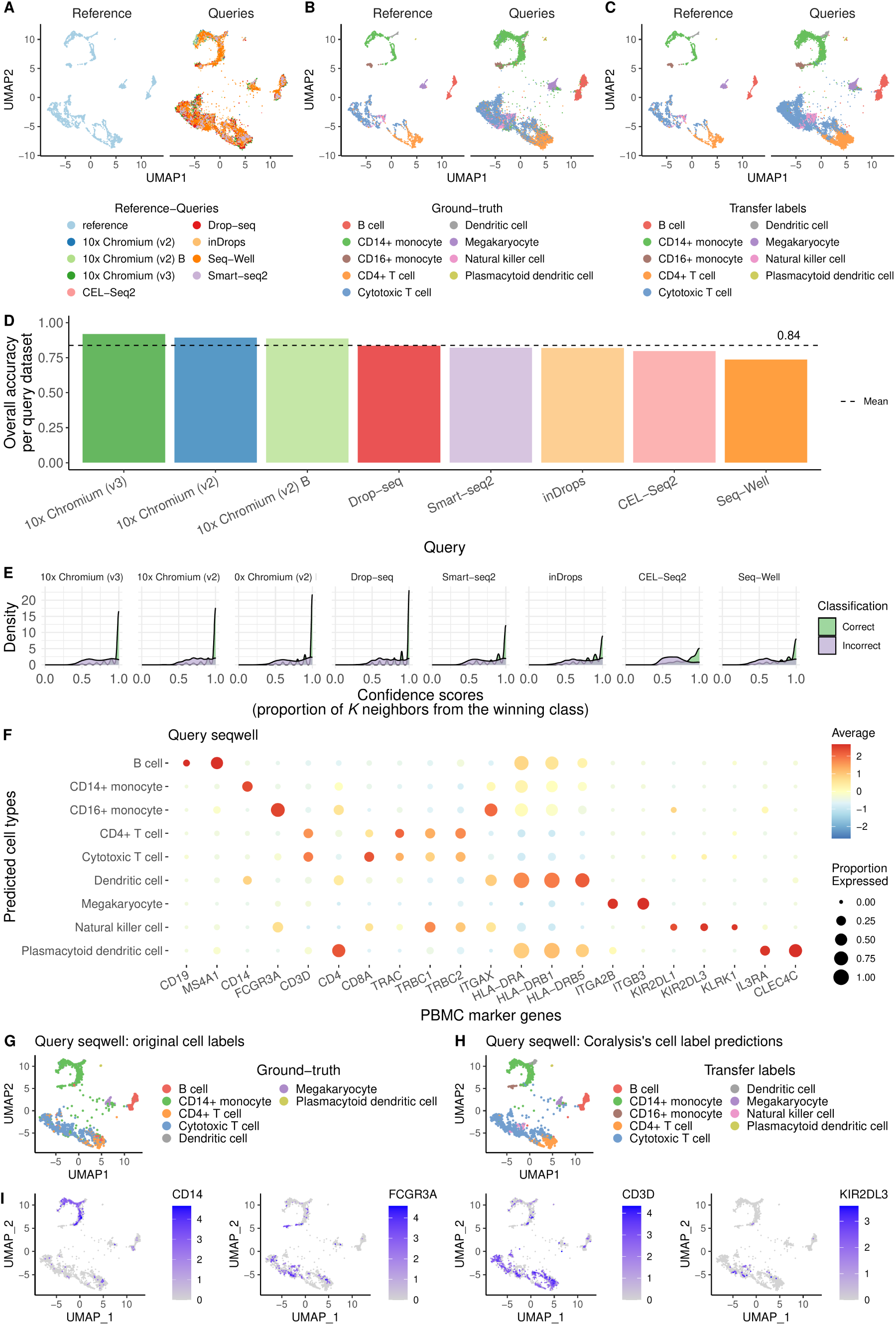
Coralysis identifies rare cell populations in a Seq-Well batch of peripheral blood mononuclear cells (PBMCs) mapped against a 10x Chromium (v2) A reference of PBMCs scRNA-seq dataset from Ding *et al.*^32^. A Queries (right) projected onto the reference (left) UMAP. Queries consisted of eight scRNA-seq datasets of PBMCs sequenced with different technologies. The reference highlighted in light blue corresponded to a PBMC scRNA-seq data prepared with 10x Chromium (v2) A technology. **B** Reference UMAP (left) and projected queries (right) highlighting the ground-truth cell type labels as distributed through *SeuratData* software from Ding *et al.*^32^. **C** Reference UMAP (left) and projected queries (right) highlighting the cell type labels transferred from the reference to the queries with Coralysis reference-mapping method. **D** Bar plot of overall accuracy per query dataset. The average accuracy across the eight queries is delimited with a dashed line. **E** Density plots of correct and incorrect classifications for each query dataset. **F** Dot plot showing the expression of marker genes for the predicted PBMC cell types for the query Seq-Well. Average expression was standardized by predicted cell type (row). Proportion expressed refers to the proportion of cells expressing the marker gene in the cell type group. UMAP of the query Seq-Well projected onto the reference highlighting **G** the ground-truth cell type labels and **H** the predicted cell labels by Coralysis. **I** UMAP of query Seq-Well showing the expression of marker genes of CD14+ monocytes (*CD14*) versus CD16+ monocytes (*FCGR3A*) as well as cytotoxic T cells (*CD3D*) versus natural killer cells (*KIR2DL3*) (from left to right).

We next inspected the reason for the lowest accuracy observed with the Seq-Well dataset. Encouragingly, all the predicted cell types were supported by the expression of their respective marker genes (Fig. 5F). Therefore, we then investigated whether the low accuracy could be due to originally wrong classifications, as the Seq-Well dataset comprised only seven cell types, while nine cell types were predicted (Fig. 5G-H). Indeed, Coralysis identified two rare populations in the dataset – CD16+ monocytes and natural killer cells – that had previously been completely missed. The identification of CD16+ monocytes, which were previously misclassified as CD14+ monocytes, was supported by the expression of *FCGR3A* (also known as *CD16*), but not *CD14*. Similarly, the appearance of natural killer cells was supported by the expression of *KIR2DL3* but not *CD3D*, a marker of cytotoxic T cells (Fig. 5I). These results demonstrated the ability of Coralysis to identify previously missed rare cell populations.

### 3.7 Coralysis cell cluster probabilities enable cell state identification

Finally, we explored the applicability of the Coralysis cell cluster probabilities for inferring cell states and their differential expression programs. The Coralysis cell cluster probability corresponds to the likelihood that a cell belongs to its assigned cluster. This probability serves as a measure of confidence of the cluster assignment and as a proxy for similarity. Therefore, we hypothesized that highly distinct cells, representing terminally differentiated or steady states, would exhibit higher probabilities, while transient or intermediate cells would show lower probabilities, as they represent cells transitioning between steady states.

To test our hypothesis, we ran Coralysis on differentiating human bone marrow CD34+ cells from Persad *et al.*^34^ (Fig. 6A). In agreement with our expectations, the Coralysis cell cluster probabilities were higher for terminally differentiated cells, such as erythroid, lymphoid, dendritic, and monocyte cell lineages, and lower for transient, fast-transitioning cells like hematopoietic stem cells and hematopoietic multipotent progenitor cells (Fig. 6B). This observation was supported by the positive Pearson correlation (*ρ*=0.73) between Coralysis cell cluster probability and Palantir pseudotime (Fig. 6C). Next, we integrated 31,029 embryonic stem cells from a developing human embryoid body, generated by Moon *et al.*^36^, with Coralysis to study the local distribution of cell cluster probabilities throughout the differentiation process (Fig. 6D,E; annotations used as given in Weiler *et al.*^37^). As expected, terminally differentiated cells such as neural crest (NC), endoderm-1 (EN-1), cardiac precursors (CPs), neuronal subtype-1 (NS-1), or hemangioblast, had higher cell cluster probabilities (Fig. 6F). Notably, Coralysis preserved inter-stage transcriptomic differences, as exemplified by embryonic stem cells and neural crest cells, despite integrating cells by embryonic stage (Fig. 6D,E). This further highlighted the ability of Coralysis to identify similar cell types across unbalanced datasets while preserving local biological differences.

**Figure 6.**
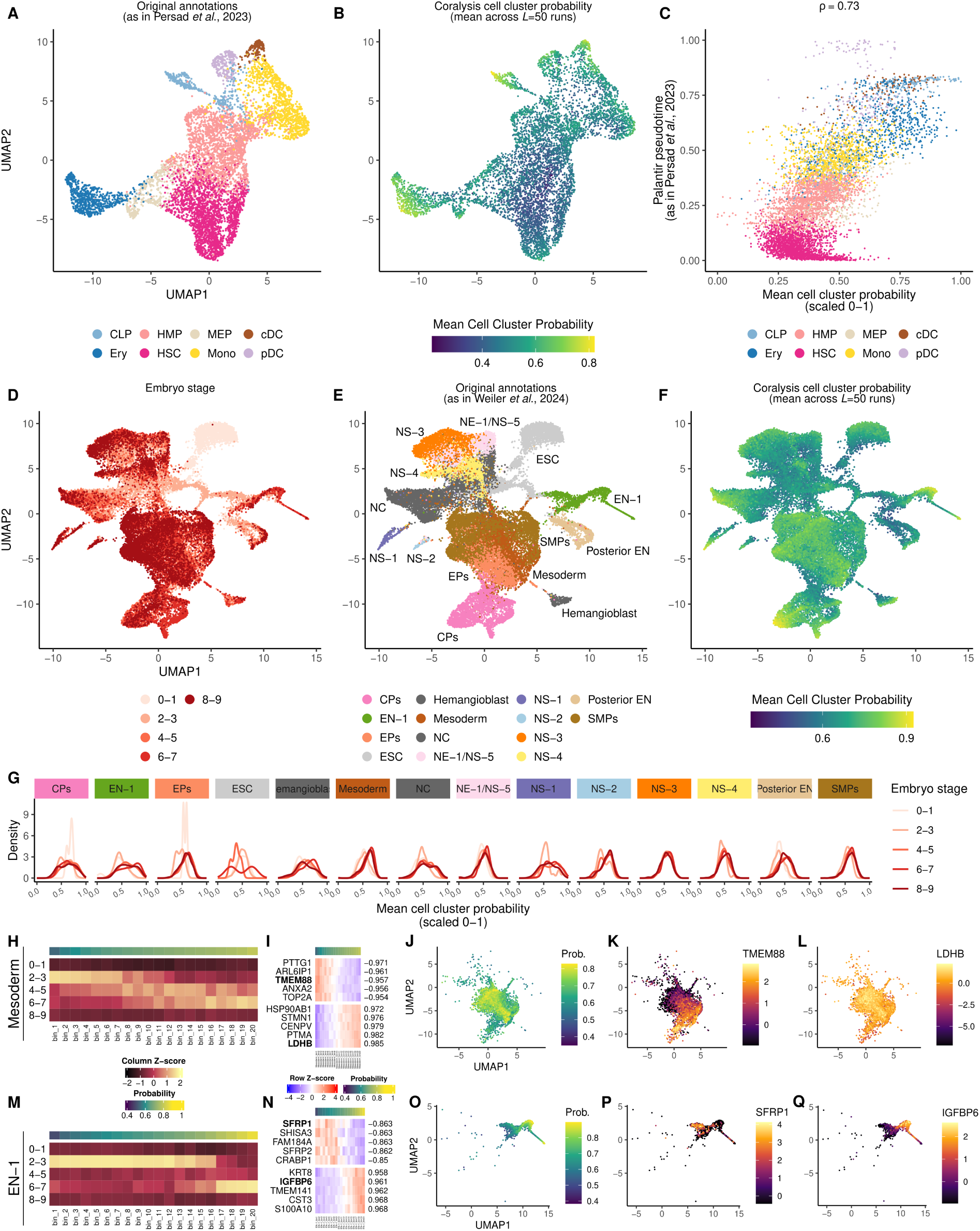
Coralysis cell cluster probability enables the identification of terminally differentiated cell types. UMAP projection of CD34+ bone marrow cells highlighting **A** cell type labels as given originally by Persad *et al.*^34^ (CLP (Common Lymphoid Progenitor), HMP (Hematopoietic Multipotent Progenitor), MEP (Megakaryocyte-Erythroid Progenitor), cDC (conventional Dendritic Cells), Ery (Erythroid), HSC (Hematopoietic Stem Cells), Mono (Monocytes), pDC (plasmacytoid Dendritic Cells)) and **B** Coralysis cell cluster probability. Mean probability across the 50 independent ICP runs (i.e., *L*=50). **C** Positive correlation between Coralysis cell cluster probability and Palantir pseudotime (*ρ*=0.73) as provided in Persad *et al.*^34^. UMAP projection of embryoid body development integrated with Coralysis highlighting **D** the embryonic stage, **E** the cell type labels as given originally by Weiler *et al.*^37^ (CPs (Cardiac Precursors), EN (Endoderm), EPs (Epicardial Precursors), ESC (Embryonic Stem Cell), Hemangioblast, Mesoderm, NC (Neural Crest), NE (Neuroectoderm), NS (Neuronal Subtypes), Posterior EN (Posterior Endoderm), SMPs (Smooth Muscle Precursors)) and **F** the Coralysis cell cluster probability. **G** Density plots of Coralysis cell cluster probability per embryonic stage for each cell type. Mean cell cluster probability was scaled to range 0-1. **H** Distribution of mesoderm cells across embryonic stages and Coralysis cell cluster probability bins (*n*=20). In total cell cluster probability was divided into 20 bins highlighted on top of the heatmap. Cell numbers were transformed by column Z-score. **I** Expression of the top five negatively and positively correlated genes across the Coralysis cell cluster probability bins for mesoderm cells. Pearson correlation was calculated between the mean cell cluster probability and the average gene expression across the 20 bins. Top colour bar corresponds to the mean cell cluster probability per bin. The averaged gene expression is represented by Z-scores (by row). UMAP representation of mesoderm cells showing **J** cell cluster probabilities and gene expression (scaled by Z-score) of **K** *TMEM88* and **L** *LDHB*. **M** Distribution of EN-1 cells across embryonic stages and Coralysis cell cluster probability bins (*n*=20). **N** Expression of the top five negatively and positively correlated genes across the Coralysis cell cluster probability bins for EN-1 cells. UMAP representation of mesoderm cells showing **O** cell cluster probabilities and gene expression (scaled by Z-score) of **P** *SFRP1* and **Q** *IGFBP6*.

Interestingly, Coralysis cell cluster probabilities were able to discriminate within-cell type differences across embryonic stages for several cell types, with cells progressing from low to high probabilities as they transitioned from early to late developmental stages (Fig. 6G).

Overall, these observations supported the applicability of Coralysis cell cluster probabilities for inferring cell states and their differential gene expression programs, which are expected to be stage-dependent, with early-stage cells being more immature and proliferative compared to mature cells at later stages. To demonstrate this, we divided the Coralysis cell cluster probabilities into 20 evenly sized bins separately for each cell type and calculated the Pearson correlation between the average cell cluster probability and the average gene expression across the bins to identify stage-specific genes. Among the top five negatively (early) and positively (late) correlated genes per cell type were genes related to cell cycle (e.g., *CDK1*, *CDKN3*), proliferation (e.g., *MALAT1*, *PTMA*), migration (e.g., *S100A10*), and differentiation (e.g., *CKB*, *ANXA2/5*) (see Supplementary Fig. 14). For instance, mesoderm and endoderm (EN-1) cells showed a good correlation between early to late developmental stages with cell cluster probability bins allowing for a better resolution over their differential gene expression programs (Fig. 6H,M). Among the top early genes for mesoderm cells, the transmembrane protein 88 (*TMEM88*) is known to inhibit the Wnt/*β*-catenin signaling pathway in order to allow the commitment of pre-cardiac mesoderm cells into cardiomyocytes^38^ (Fig. 6I-K). Notably, this was supported by the co-localisation of *TMEM88*-expressing cells with epicardial and cardiac precursors (EPs and CPs; Fig. 6E). Additionally, glycolysis-related genes, such as *LDHB*, which was upregulated in late mesoderm cells, have shown differential expression across developmental stages in mesoderm during mouse gastrulation^39^ (Fig. 6L). Finally, in agreement with our findings, Vianello & Lutolf^40^ found *SFRP1* and *IGFBP5*, the latter functionally similar to *IGFBP6*, as being upregulated in early and late endoderm, respectively (Fig. 6N-Q). Overall, these results highlighted the ability of Coralysis to identify functionally relevant cell states and their differential expression programs.

## 4 Discussion

In this study, we present Coralysis, a method for sensitive integration, reference-mapping, and cell state identification across single-cell datasets through multi-level integration. It relies on an adapted version of our previously introduced Iterative Clustering Projection^15^ (ICP) algorithm to identify shared cell clusters across heterogeneous datasets by leveraging multiple rounds of divisive clustering. Similar to assembling a “puzzle”, multi-level integration enables ICP to blend the batch effects and separate cell types by focusing on low- to high-level features (see Fig. 1 and Extended Data Fig. 2). The trained ICP models can then be used for various purposes, including prediction of cluster identities of related, unannotated single-cell datasets through reference-mapping, and inference of cell states and their differential expression programs using the cell cluster probabilities that represent the likelihood of each cell belonging to each cluster. Coralysis performed consistently well across a wide range of integration tasks compared to current top-performing integration methods. It prioritises bio-conservation over batch correction, outperforming existing methods in integration tasks involving datasets that do not share similar cell-types. Furthermore, Coralysis exhibited minimal variation in performance based on input data type, partly due to its inherent feature extraction procedure, which employs *L1*-regularized logistic regression^15^.

The feature extraction procedure of Coralysis is particularly relevant for the detection of diseased or perturbed cell states and rare populations, as feature selection methods might overlook features required to discriminate these. For example, the commonly used Seurat highly variable gene selection method relies largely on lowly expressed genes with high coefficients of variation representing technical noise that masks subtle cellular variation^41^. Furthermore, Coralysis allows the retrieval of gene coefficients from ICP models for a given ICP run or clustering label using a majority voting approach. This offers the advantage of identifying cluster marker genes independently from differential expression analysis avoiding the bias towards highly expressed genes representing false-positives^42,43^. Notably, logistic regression, which is at the core of ICP, was one of the best methods for identifying cluster markers among 59 recently benchmarked tools^44^.

As the complexity of cell atlases increases in terms of tissue, time, and condition representation, subtle biological differences can become difficult to dissociate from batch effects, likely increasing the demand for such integration functionalities. The application of Coralysis is not limited to single-cell transcriptomics; it can also be applied to integrate data from other single-cell assays, such as antibody-derived tags (ADT) or cytometry by time of flight (CyTOF), within the exciting and rapidly evolving field of single-cell proteomics (Fig. 3). The reduced feature space of these assays, typically ranging from dozens to a few hundred, did not prevent Coralysis from identifying fine-grained cell populations and preserving differentiation trajectories, nor from detecting rare populations, such as basophils (0.5%), in whole blood CyTOF data. The identification of cell populations at high resolution provided by Coralysis is expected to be of great value as unbiased measurements of single-cell proteomes become more common with advancements in mass spectrometry^45^. In the future, we aim to extend the application of the Coralysis integration method to other single-cell assays, such as ATAC-seq, and even across different modalities with shared features by testing different regularizations (e.g., Ridge instead of LASSO) and data input transformations.

Coralysis also allows to avoid cumbersome *de novo* integration when a reference trained with Coralysis exists. Reference-mapping is becoming routine as projects like the Human Cell Atlas^46^ or the Tabula Muris^47^ work to decipher the identities of millions of cells. Robust methods are needed to accurately map, for instance, diseased cells against healthy references, perturbed cells, or cells generated by new single-cell technologies. Here, we demonstrated that Coralysis performs well across several query-reference scenarios, including cell-type imbalance, batch representation, or varied batch strength. We also showed the applicability of Coralysis to identify previously missed rare cell populations across query-reference datasets originating from different sequencing technologies. Additionally, the confidence scores provided help for the user to identify which cell type predictions should be more carefully inspected. Although Coralysis currently has limitations in detecting cells not represented in the reference, this is unlikely to pose a hurdle as references become increasingly comprehensive. Future developments should address this limitation, for instance, by comparing the similarity between query and reference model coefficients.

A long-standing challenge in single-cell biology is the definition of cell type versus cell state and how to identify and distinguish them. Cell states correspond to discrete modes that a given cell type can assume under different stimuli^48^. As such, they share a common gene expression program, which places them under the same cell type ‘umbrella’ definition. For example, the CD8+ T cell type can assume the naive, memory, or effector CD8+ T cell states. Coralysis cell cluster probability aids the identification of terminally differentiated and transient cell types. Similarly to ‘metacells’, cell cluster probability bins aggregate highly similar cells, representing primarily technical noise, and captures existing biological variation within cell types across bins. This can help unravel cell states and their differential expression programs, such as those involved in early to late embryoid cell development or healthy versus diseased cells. We anticipate that Coralysis will advance the catalog of cell heterogeneity by improving the integration of imbalanced cell types and states, enabling a more faithful representation of the cellular landscape in complex single-cell experiments.

## Acknowledgements

The authors thank the Elo lab for useful discussions and support. AGGS and LLE were supported by the European Union’s Horizon 2020 research and innovation programme under the Marie Skłodowska-Curie grant agreement no.: 955321. AGGS was also supported by the University of Turku, Å bo Akademi University, Turku Graduate School (UTUGS). LLE reports grants from Research Council of Finland (310561, 329278, 335434, 335611, 341342, 364700), Sigrid Juselius Foundation, and Cancer Foundation Finland during the conduct of the study. Our research is also supported by University of Turku Graduate School (UTUGS), Biocenter Finland, and ELIXIR Finland.

## 5 Author contributions

AGGS conceptualised and implemented the method, performed the analyses, prepared the figures and wrote the manuscript. JS assisted in the conceptualisation and analyses. SJ assisted in the conceptualisation and analyses, tested the method, critically reviewed the analyses and supervised the work. LLE conceived and supervised the study, tested the method, critically reviewed the analyses, and participated in writing the manuscript. All authors reviewed and approved the final version of the manuscript.

---

**Extended Data Figure 1.**
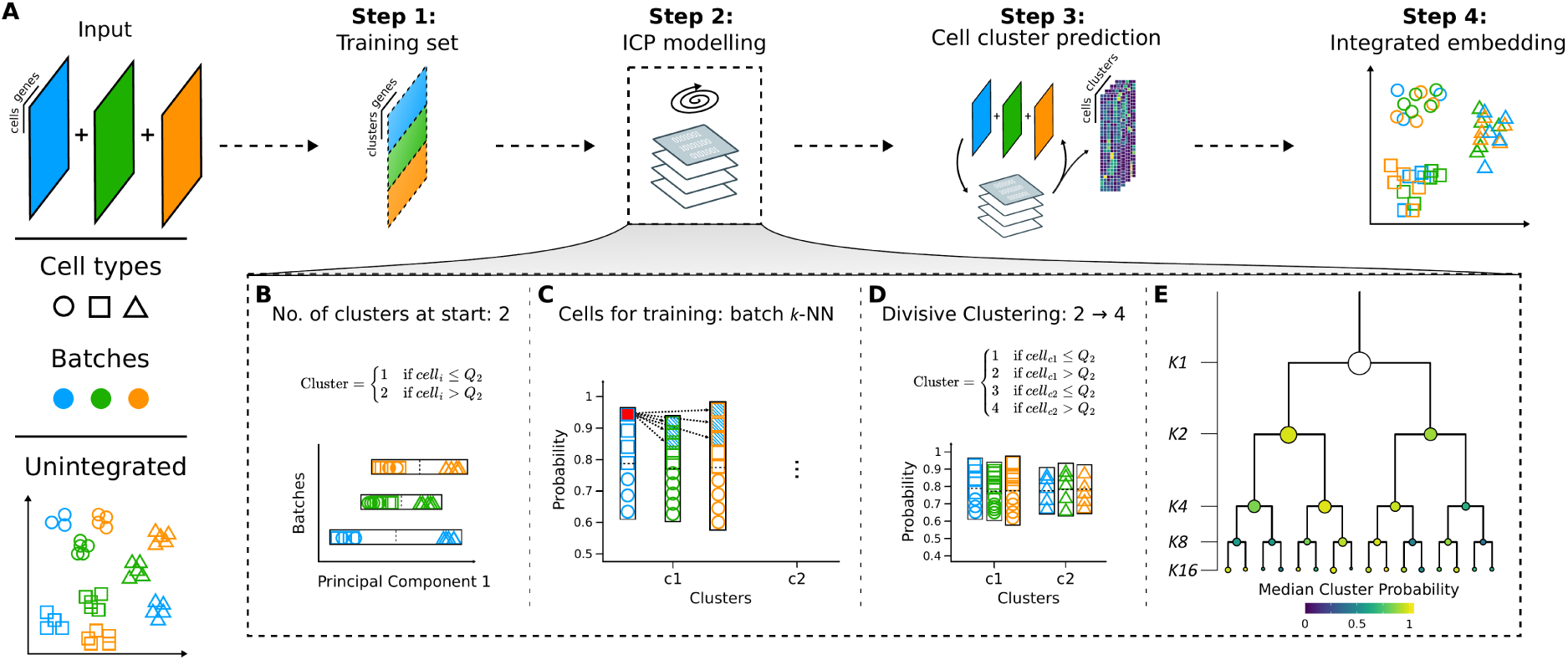
Coralysis integration flowchart. **A** An input of heterogeneous scRNA-seq datasets are overclustered batch wise into a training set modelled through the Iterative Clustering Projection (ICP) algorithm in order to predict the cell cluster probabilities and obtain an integrated embedding. Adaptations to the original ICP algorithm^15^: **B** batch wise cluster assignment at start, dependent on the cell distribution across Principal Component 1 (median as cutoff); **C** training cells selected from batch *k* nearest neighbours of the cell with the highest probability for every batch per cluster; and, **D** upon ICP clustering convergence, each cluster is further divided into two for the next clustering round, dependent on the batch wise cluster probability distribution (median as cutoff). **E** Multi-level integration is achieved through multiple divisive clustering rounds, blending the batch effect and highlighting the biological signal incrementally. Shapes represent cell types and colours batches.

**Extended Data Figure 2.**
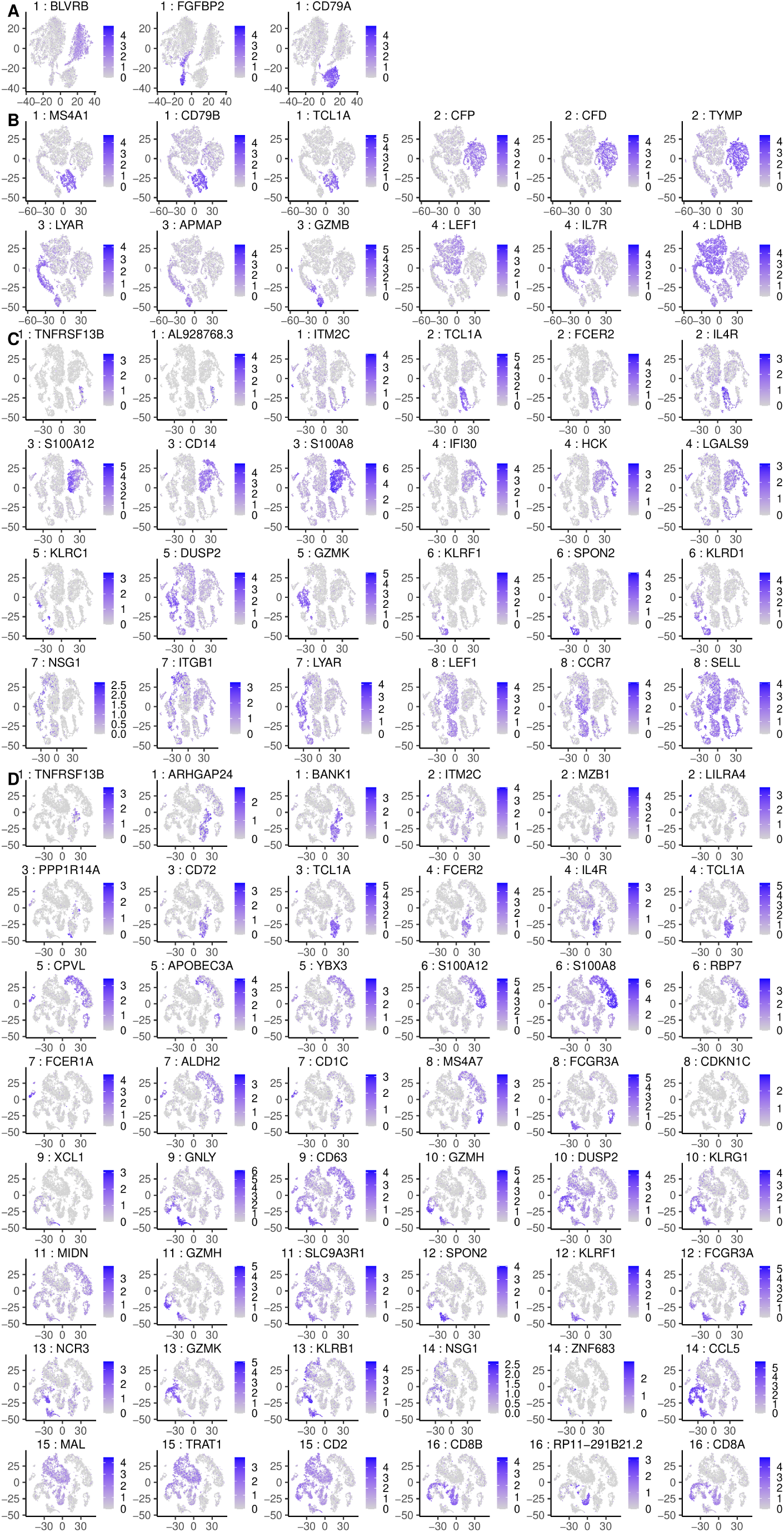
Expression of the top three positive gene coefficients for the ICP model corresponding to the run *L=2* projected onto *t* -SNE. A-D. Top three positive coefficients per cluster per clustering round level *K2* **A**, *K4* **B**, *K8* **C** and *K16* **D**. The plot title highlights the cluster number and the respective gene coefficient. For every clustering round level a *t* -SNE was built from the concatenation of all cluster probability tables (*L*=50 ICP runs) from the respective clustering round.

**Extended Data Figure 3.**
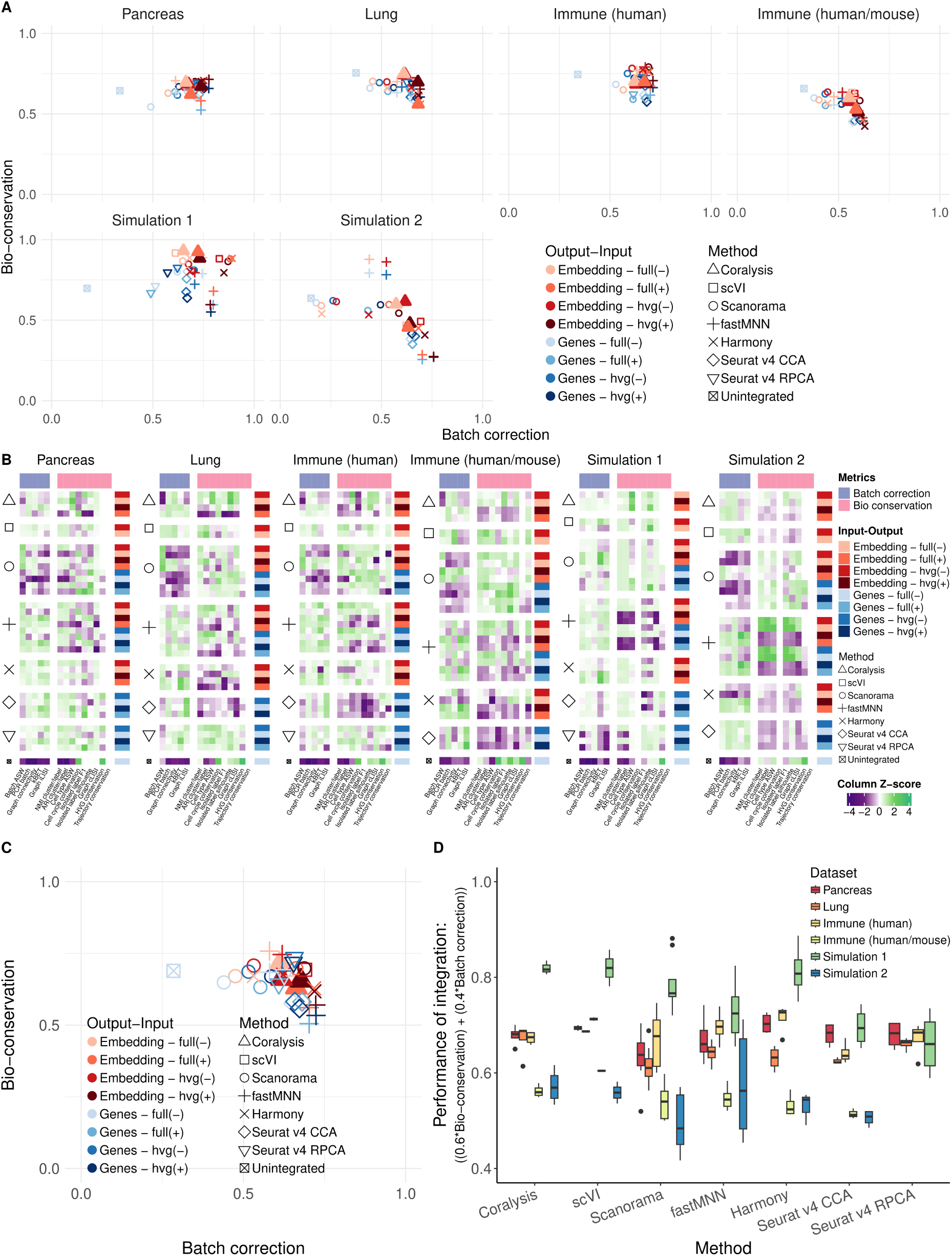
Benchmark of Coralysis integration method through the scib-pipeline. **A** Mean of batch-correction (*n*=5) versus bio-conservation (*n*=9) scib metrics for each method benchmarked (Coralysis, scVI, Scanorama, fastMNN, Harmony, Seurat v4 CCA/RPCA) across four real (pancreas, lung, human immune, human/mouse immune) and two simulated datasets (simulation 1 and 2). Colours represent different output-input and shapes distinct methods. Different types of output consist of an integrated embedding (embedding) or a batch-corrected gene expression matrix (genes). The different inputs included providing the gene expression data scaled (+) or unscaled (-) and with (hvg) or without (full) feature selection. **B** Individual bio-conservation and batch-correction scib metrics for each method and dataset benchmarked. Metrics were standardized (Z-score) across methods. **C** Averaged bio-conservation and batch-correction scores across the six datasets. **D** Variance in performance of integration for different input-output(s) across methods. Performance score consists of the weighted average of bio-conservation (0.6) and batch-correction (0.4) metrics.

**Extended Data Figure 4.**
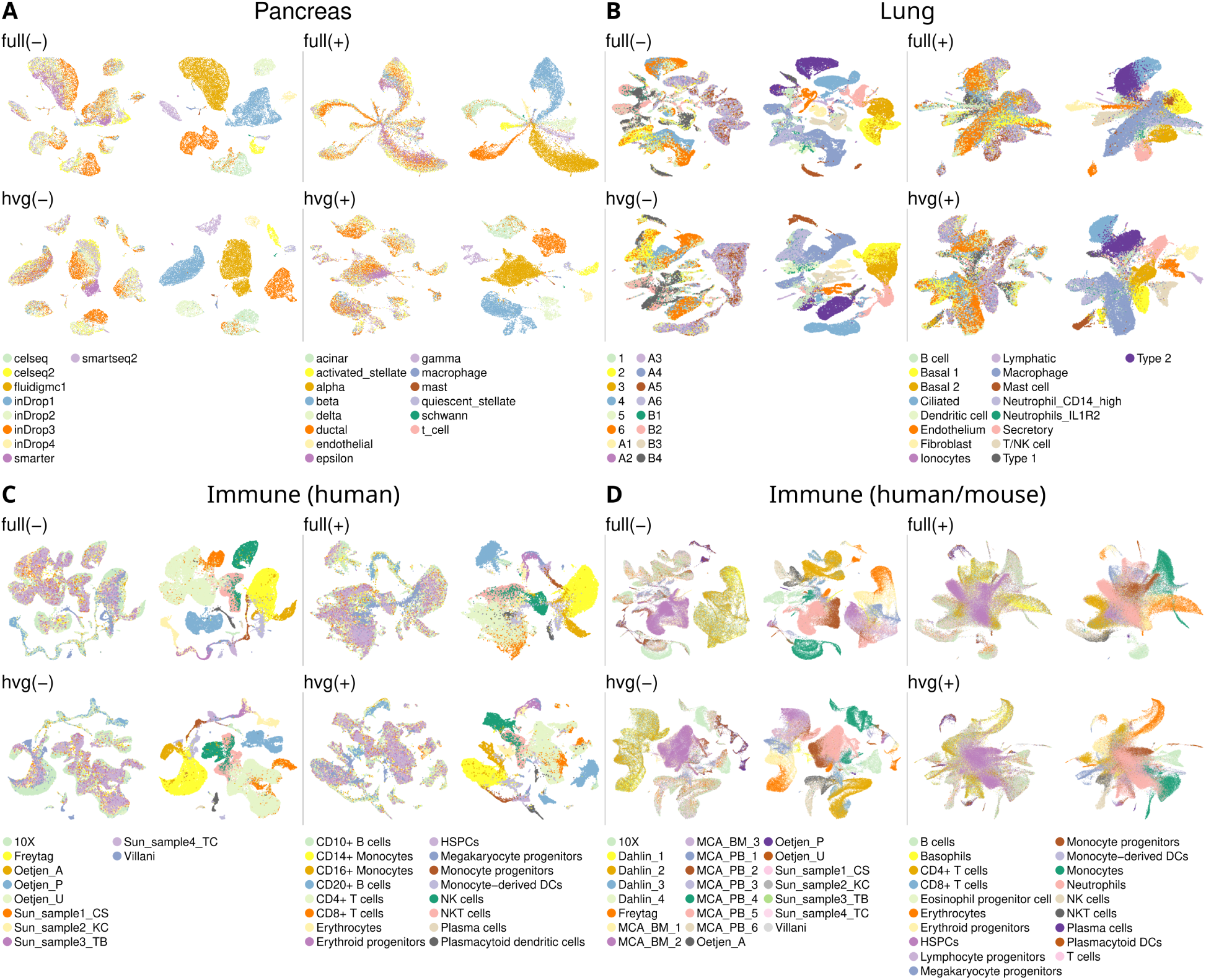
Coralysis integrated UMAP projections for the four real datasets obtained through the scib-pipeline benchmark. The datasets included were pancreas, lung atlas, human immune and human/mouse immune **A-D**. The top left subtitle corresponds to the input data used (from left to right, from top to bottom): full unscaled gene expression matrix (full(-)), full scaled gene expression matrix (full(+)), highly variable genes unscaled gene expression matrix (hvg(-)) and highly variable genes scaled gene expression matrix (hvg(+)). The left and right plots for each dataset for a given input data type highlight the batch and cell type labels, respectively.

**Extended Data Figure 5.**
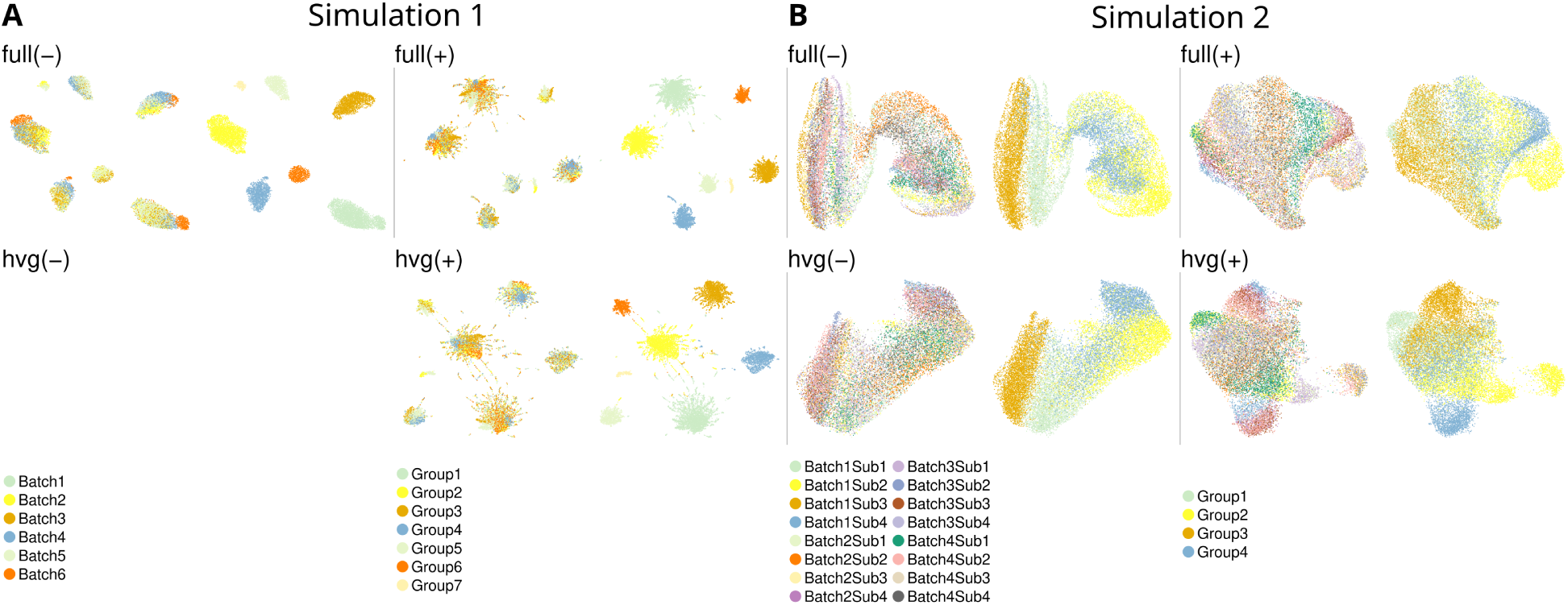
Coralysis integrated UMAP projections for the two simulated datasets obtained through the scib-pipeline benchmark. The datasets included were simulations 1 and 2 **A-B**. The top left subtitle corresponds to the input data used (from left to right, from top to bottom): full unscaled gene expression matrix (full(-)), full scaled gene expression matrix (full(+)), highly variable genes unscaled gene expression matrix (hvg(-)) and highly variable genes scaled gene expression matrix (hvg(+)). The left and right plots for each dataset for a given input data type highlight the batch and cell type labels, respectively. The integration task for the simulation 1 dataset, given the input hvg(-), failed.

**Extended Data Figure 6.**
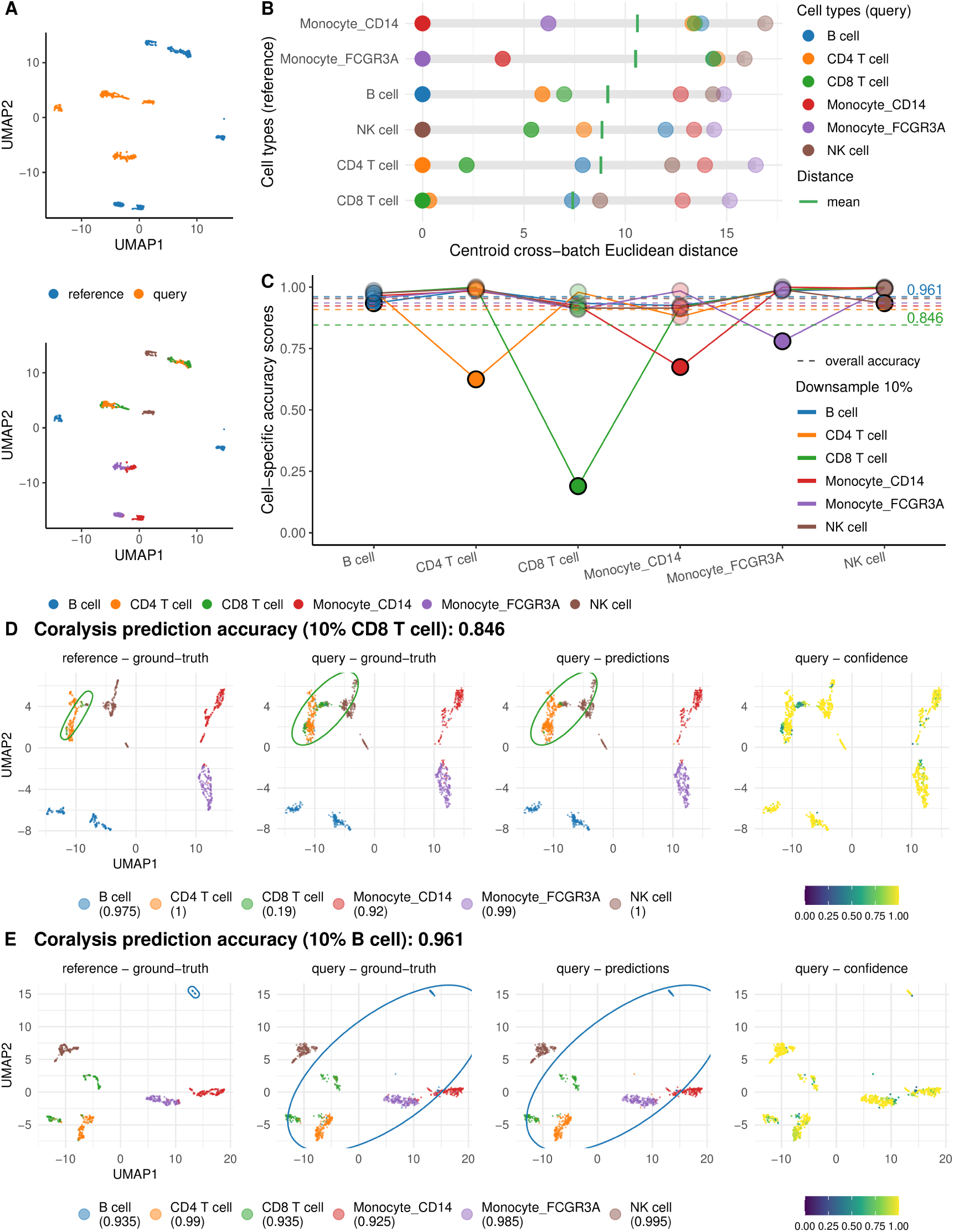
Accuracy performance of Coralysis reference-mapping method for a query-reference scenario of imbalanced cell types. **A** UMAP projection of query and reference PBMC datasets coloured by dataset (top) and cell type (bottom) identities. **B** Centroid cross-batch Euclidean distance in PCA space highlighting the similarity between cell types across batches. The distance between the same cell type across reference-query for every reference-query comparison was subtracted in the respective comparison in order to set the distance of every comparison to start from zero. **C** Cell-specific accuracy scores in down-sampled comparisons. In each reference-query comparison one cell type was down-sampled to 10%. The accuracy scores for each cell type in the same comparison are joined by the solid lines, their colours representing the down-sampled cell type. The average accuracy score for each comparison is given by the dashed line, with the two values highlighted corresponding to the lowest and highest overall accuracy (CD8 T cell-B cell). For every comparison, the darkest dot corresponds to the down-sampled cell type. Projection of query onto reference UMAP highlighting ground-truth, predictions and confidence scores obtained with Coralysis reference-mapping method for the comparisons with the lowest **D** and the highest **E** overall accuracy (CD8 T cell-B cell). The confidence scores represent the proportion of *K* neighbours from the winning class (*K* =10). Ellipses circumscribe the position of the down-sampled cell type.

**Extended Data Figure 7.**
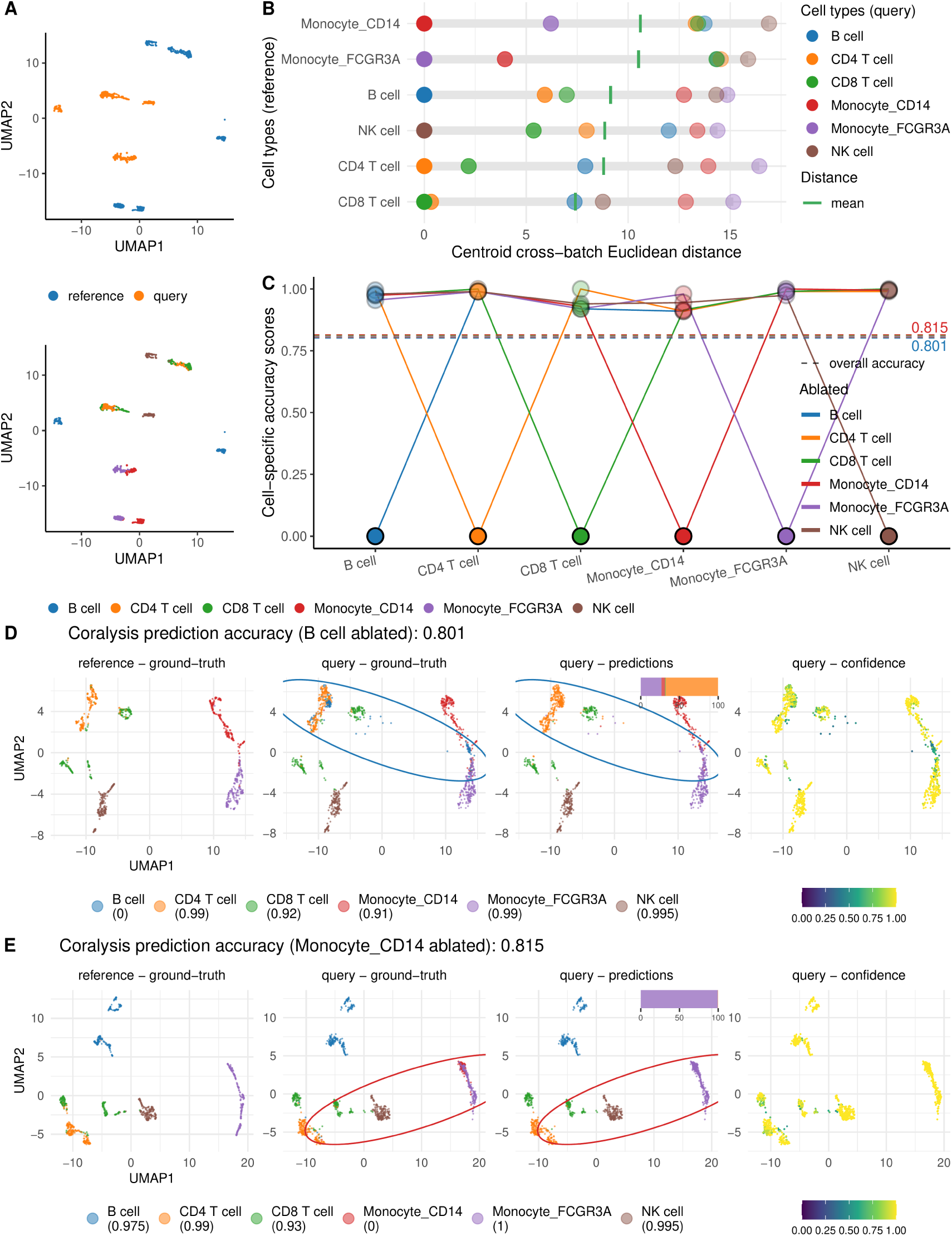
Accuracy performance of Coralysis reference-mapping method for a query-reference scenario of unshared cell types. **A** UMAP projection of query and reference PBMC datasets colored by dataset (top) and cell type (bottom) identities. **B** Centroid cross-batch Euclidean distance in PCA space highlighting the similarity between cell types across batches. The distance between the same cell type across reference-query for every reference-query comparison was subtracted in the respective comparison in order to set the distance of every comparison to start from zero. **C** Cell-specific accuray scores in ablated comparisons. In each reference-query comparison one cell type was ablated. The accuracy scores for each cell type in the same comparison are joined by the solid lines, their colours representing the ablated cell type. The average accuracy score for each comparison is given by the dashed line, with the two values highlighted corresponding to the lowest and highest overall accuracy (B cell-CD14 monocyte). For every comparison, the darkest dot corresponds to the ablated cell type. Projection of query onto reference UMAP highlighting ground-truth, predictions and confidence scores obtained with Coralysis reference-mapping method for the comparisons with the lowest **D** and the highest **E** overall accuracy (B cell-CD14 monocyte). The bar plot above the UMAP corresponds to the cell type labels against which the query cell type, ablated in the reference, was classified. The confidence scores represent the proportion of *K* neighbours from the winning class (*K* =10). Ellipses circumscribe the position of the ablated cell type.

**Extended Data Figure 8.**
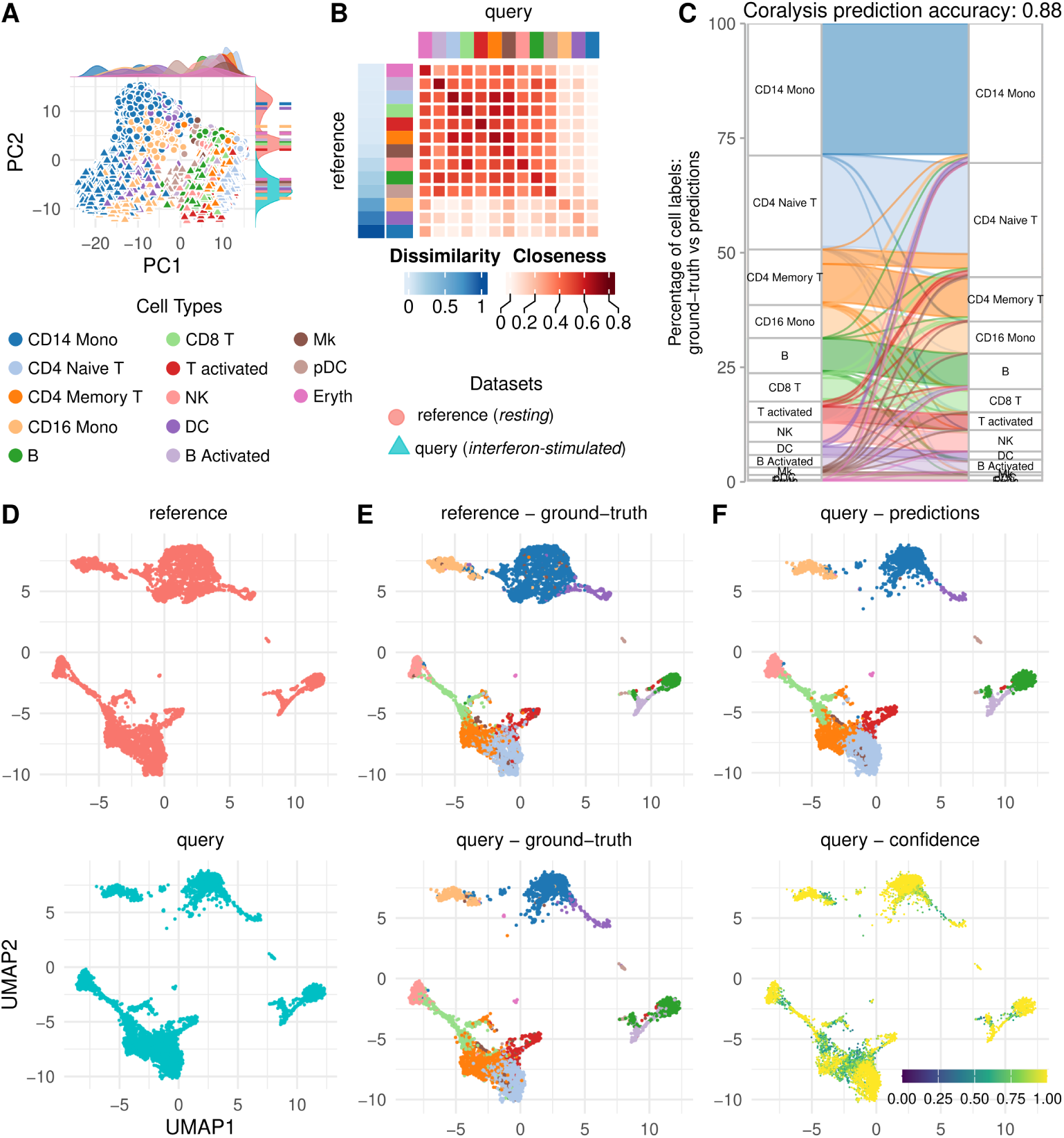
Accuracy performance of Coralysis reference-mapping method for a query-reference scenario of varied strength of the batch effect. **A** Joint PCA of reference and query PBMC datasets. Reference and query consisted of resting and interferon stimulated PBMCs. Shapes represent reference-query identity and colours cell types. Distributions on top and left highlight cell types and dataset identity across PC1 and PC2, respectively. Dashed lines across PC2 dataset-identity distributions correspond to the cell type centroids across reference-query. **B** Cross-batch centroid (Euclidean) distance between reference-query cell types. Dissimilarity corresponds to the diagonal distances min-max scaled. Closeness corresponds to the Euclidean distances scaled to fit in the range [0-1] by subtracting from one the distance divided by the maximum distance. **C** Sankey plot showing the correspondence between ground-truth cell type labels and predictions in percentage. Query projected onto the reference UMAP highlighting the dataset identity **D**, ground-truth cell labels **E**, predictions and confidence scores **F** between the reference and query (top-bottom). The confidence scores represent the proportion of *K* neighbours from the winning class (*K* =10).

