## Supplementary Material for "Coralysis enables sensitive identification of imbalanced cell types and states in single-cell data via multi-level integration"

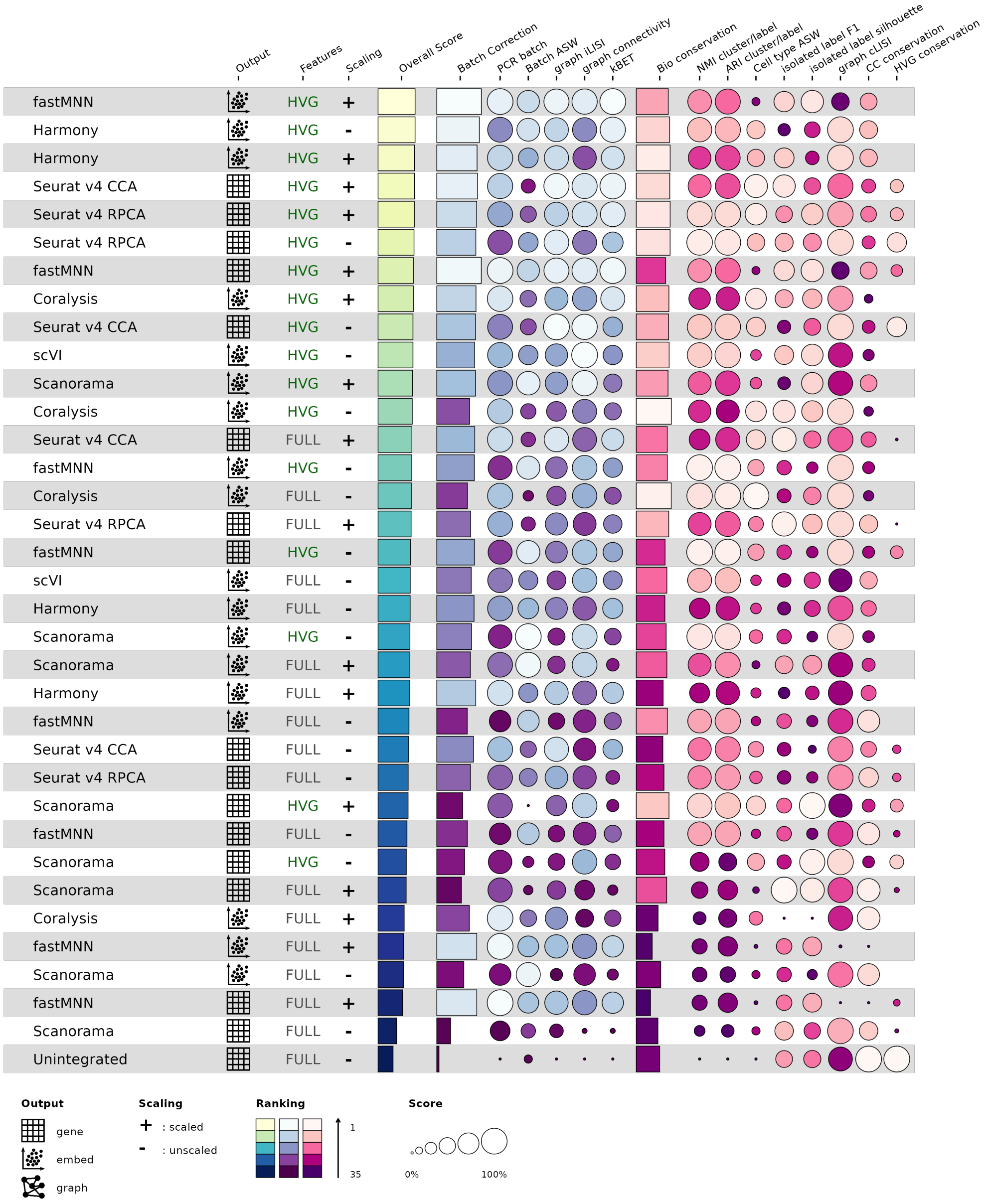


**Supplementary Figure 1 Performance ranking of integration methods by the overall score obtained with the scib-pipeline for the pancreas dataset.** Overall score corresponds to 0.4:0.6 weighted mean between batch-correction (blue/purple) and bio-conservation (pink) metrics, respectively.

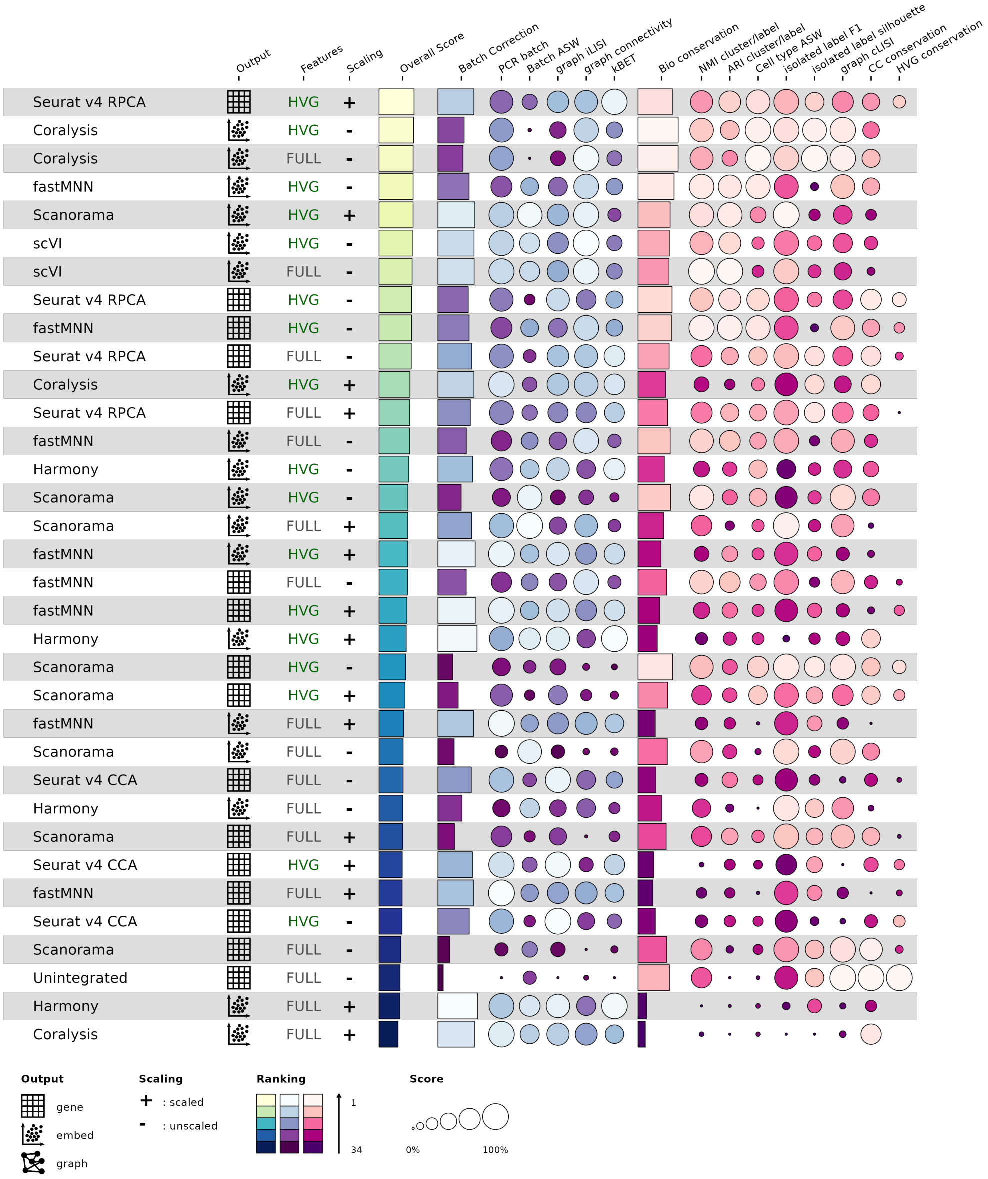


**Supplementary Figure 2 Performance ranking of integration methods by the overall score obtained with the scib-pipeline for the lung atlas dataset.** Overall score corresponds to 0.4:0.6 weighted mean between batch-correction (blue/purple) and bio-conservation (pink) metrics, respectively.

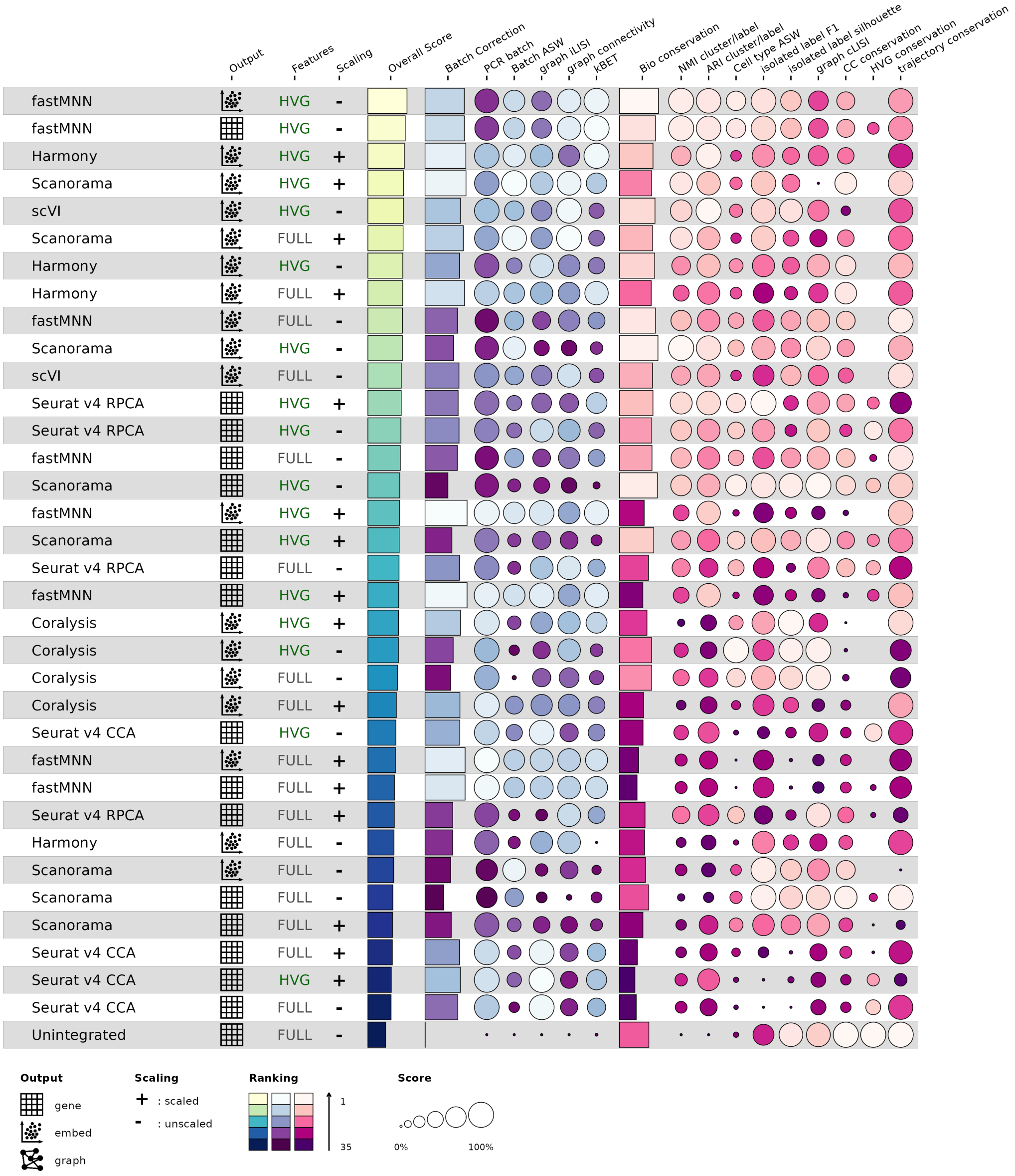


**Supplementary Figure 3 Performance ranking of integration methods by the overall score obtained with the scib-pipeline for the human immune dataset.** Overall score corresponds to 0.4:0.6 weighted mean between batch-correction (blue/purple) and bio-conservation (pink) metrics, respectively.

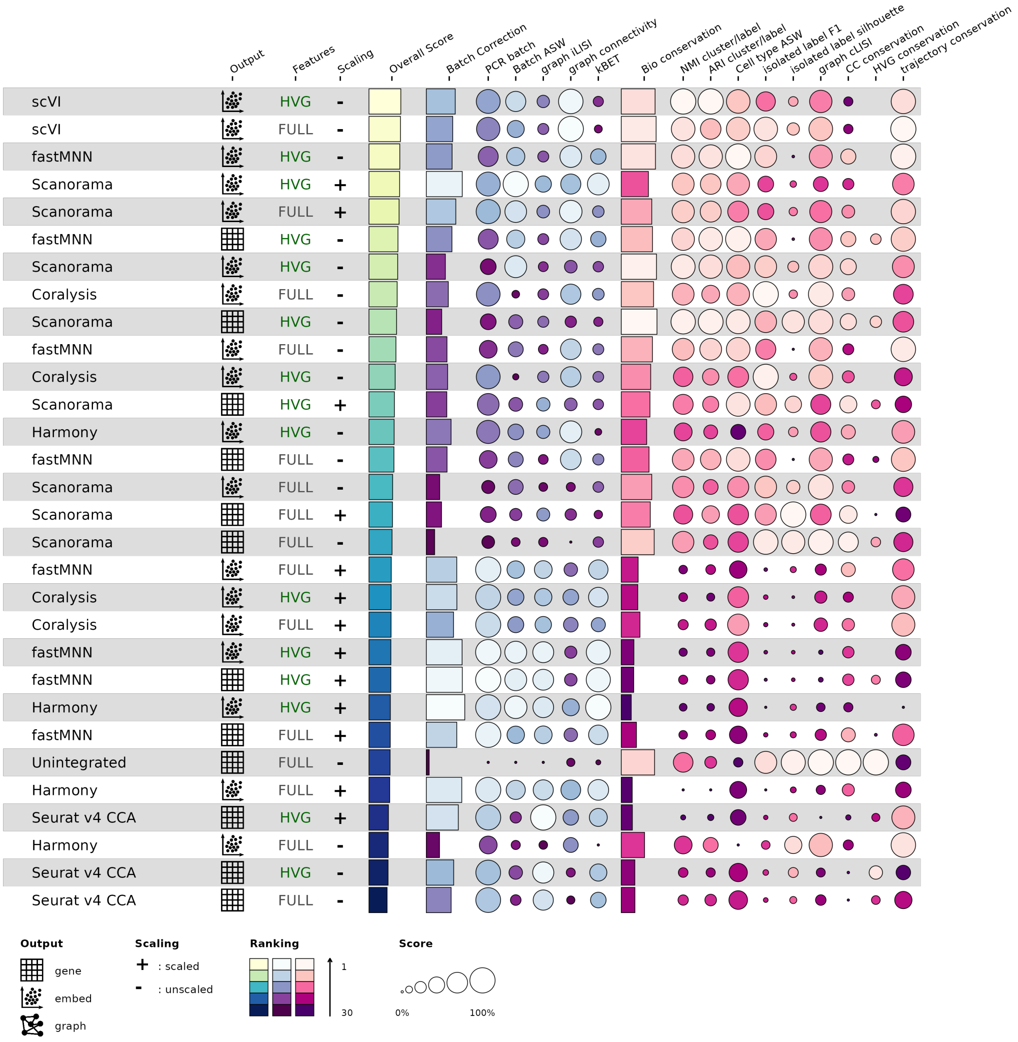


**Supplementary Figure 4 Performance ranking of integration methods by the overall score obtained with the scib-pipeline for the human/mouse dataset.** Overall score corresponds to 0.4:0.6 weighted mean between batch-correction (blue/purple) and bio-conservation (pink) metrics, respectively.

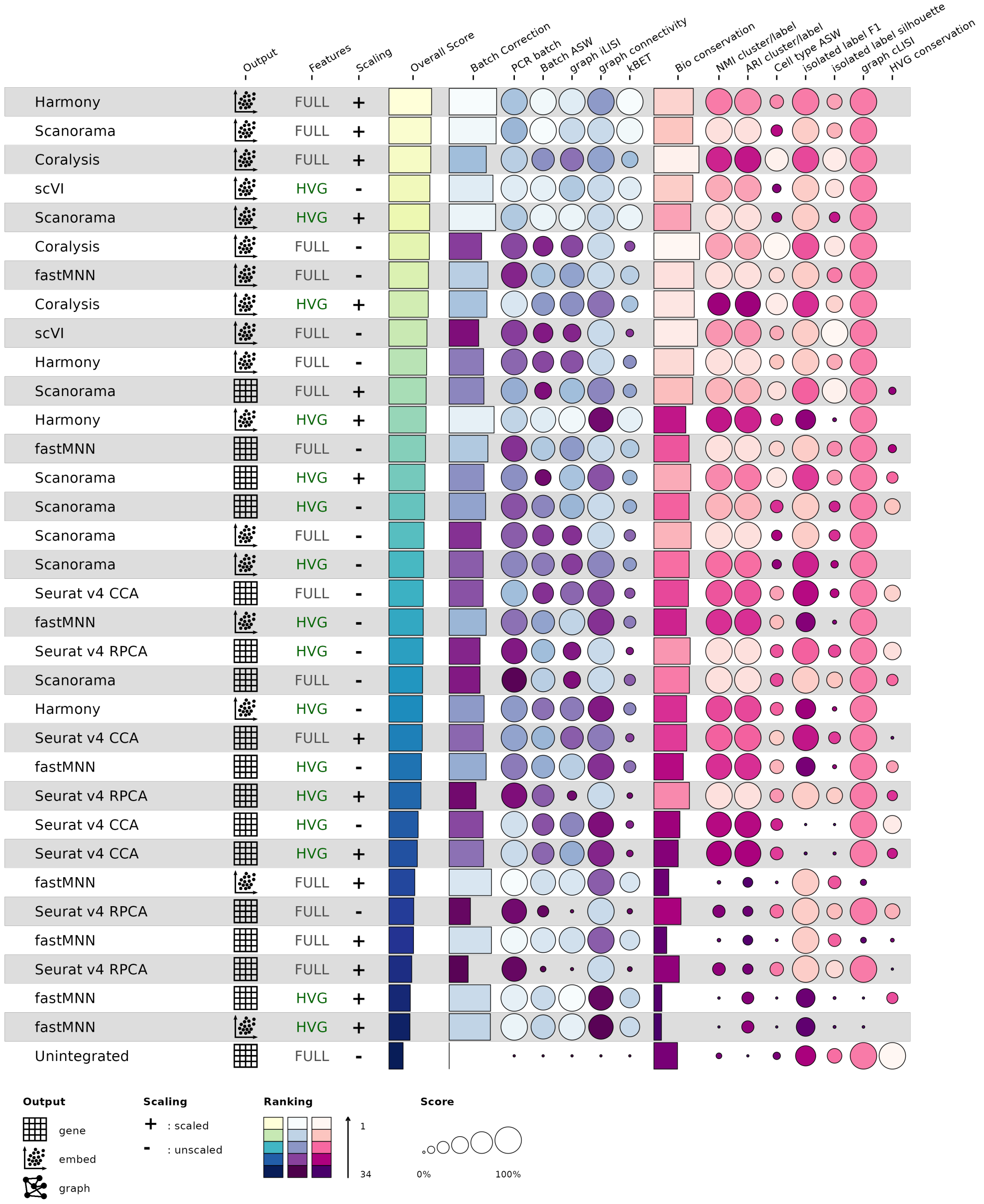


**Supplementary Figure 5 Performance ranking of integration methods by the overall score obtained with the scib-pipeline for the simulation 1 dataset.** Overall score corresponds to 0.4:0.6 weighted mean between batch-correction (blue/purple) and bio-conservation (pink) metrics, respectively.

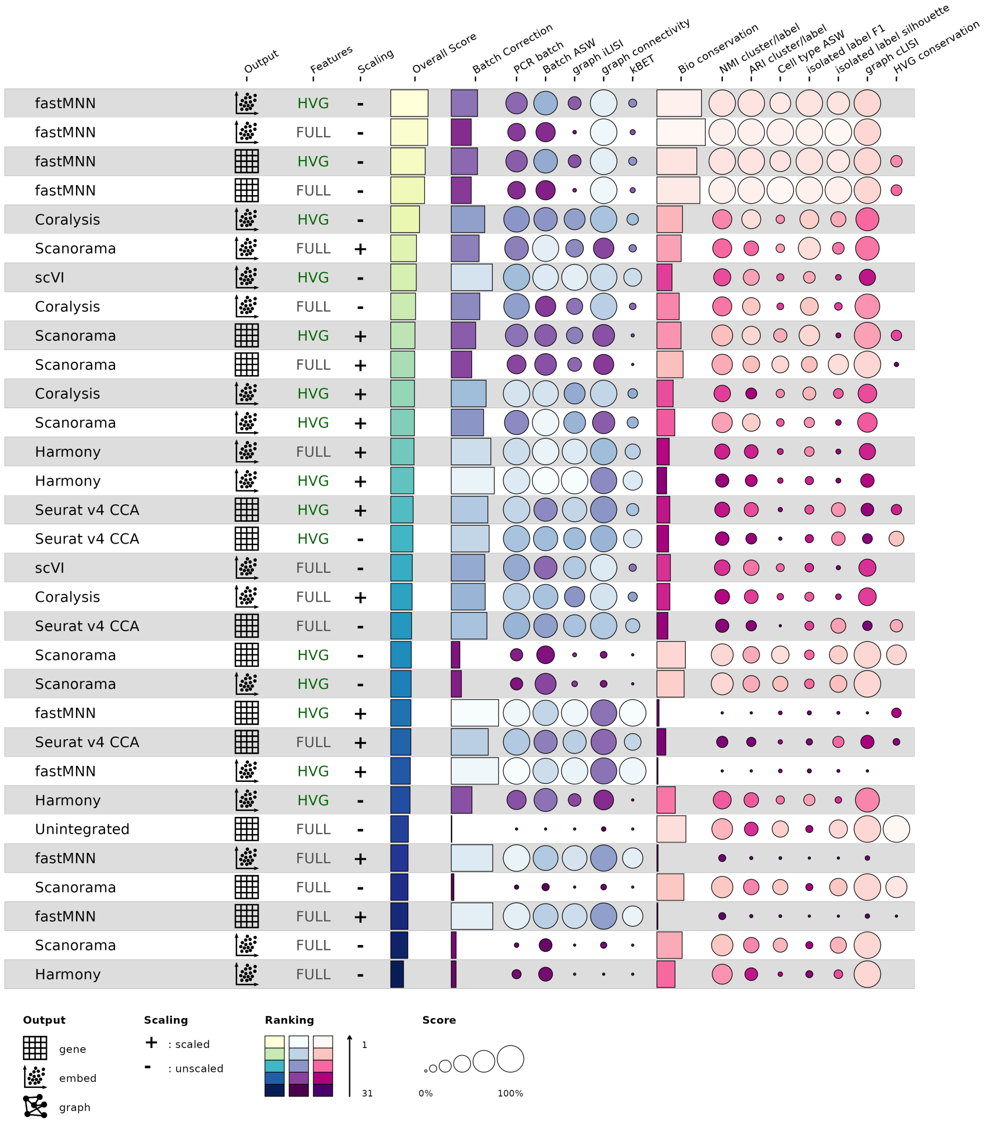


**Supplementary Figure 6 Performance ranking of integration methods by the overall score obtained with the scib-pipeline for the simulation 2 dataset.** Overall score corresponds to 0.4:0.6 weighted mean between batch-correction (blue/purple) and bio-conservation (pink) metrics, respectively.

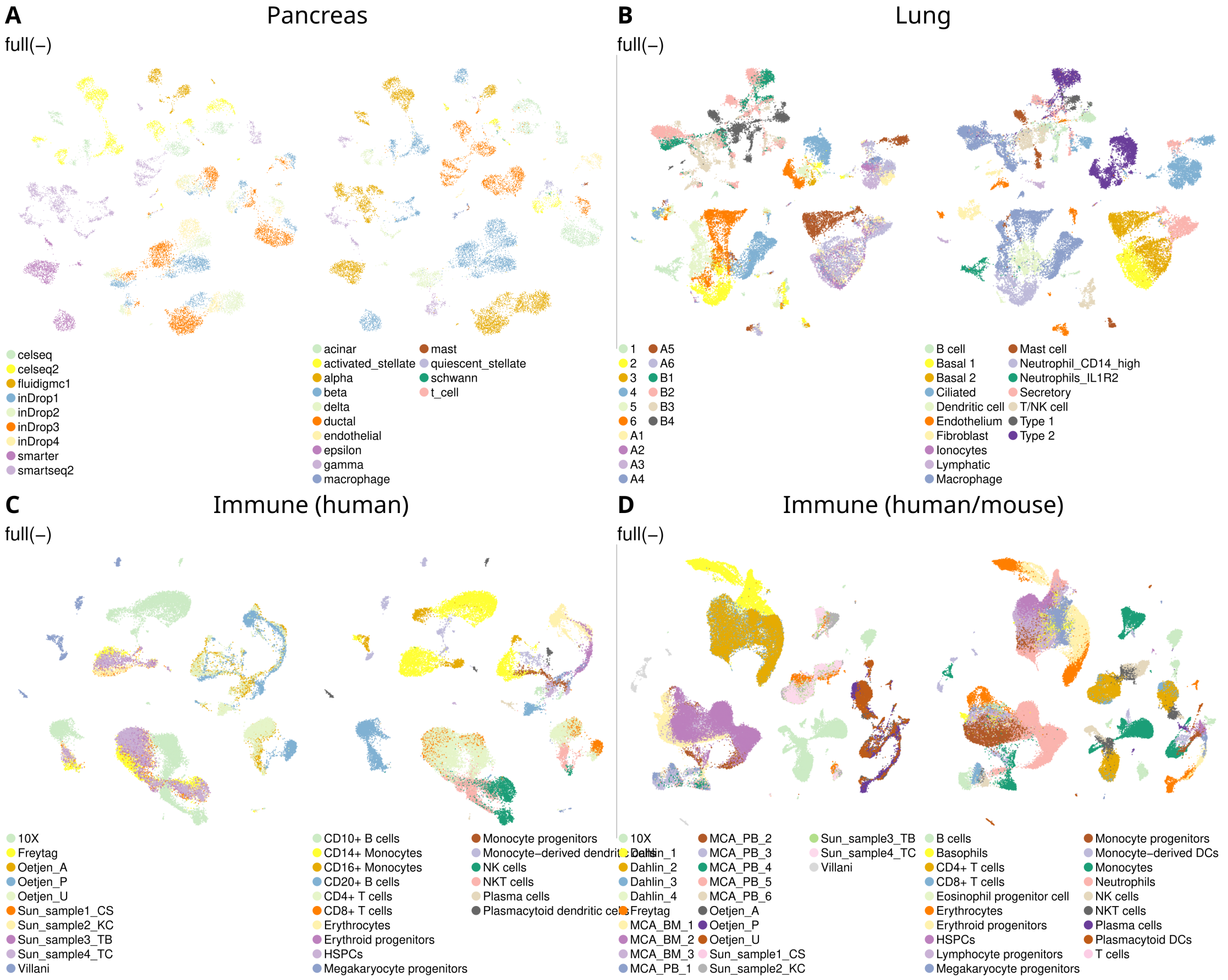


**Supplementary Figure 7 Unintegrated UMAP projections for the four real datasets used to benchmark Coralysis obtained through the scib-pipeline.** The datasets included were pancreas, lung atlas, human immune and human/mouse immune **A-D**. The centered title corresponds to the dataset. The top left subtitle corresponds to the input data used, i.e., full unscaled gene expression matrix (full(-)). The left and right plots for each dataset highlight the batch and cell-type labels, respectively.

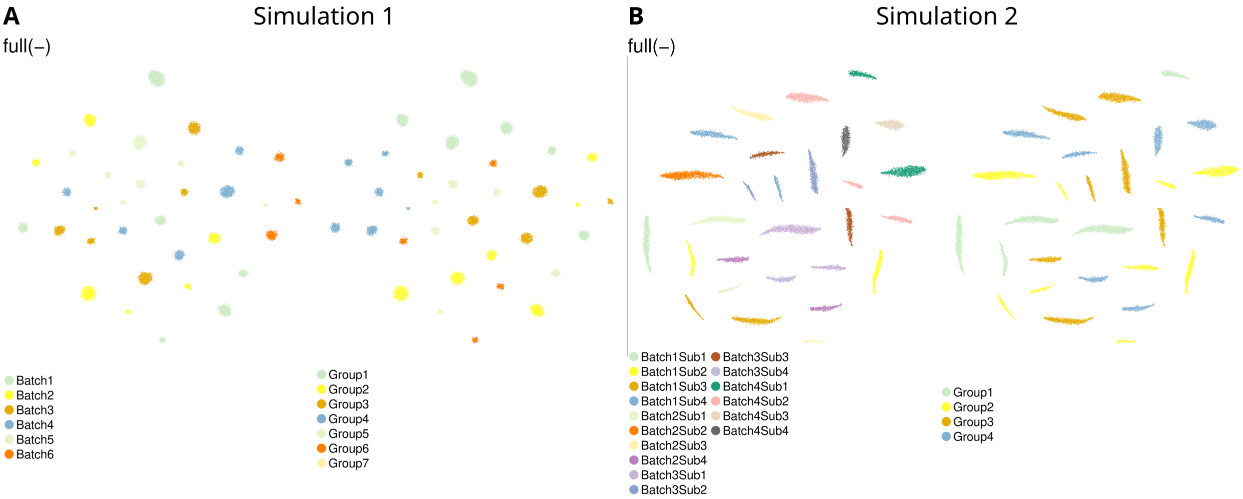


**Supplementary Figure 8 Unintegrated UMAP projections for the two simulated datasets used to benchmark Coralysis obtained through the scib-pipeline.** The datasets included were simulation 1 and 2 **A-B**. The centered title corresponds to the data set. The top left subtitle corresponds to the input data used, i.e., full unscaled gene expression matrix (full(-)). The left and right plots for each dataset highlight the batch and cell-type labels, respectively.

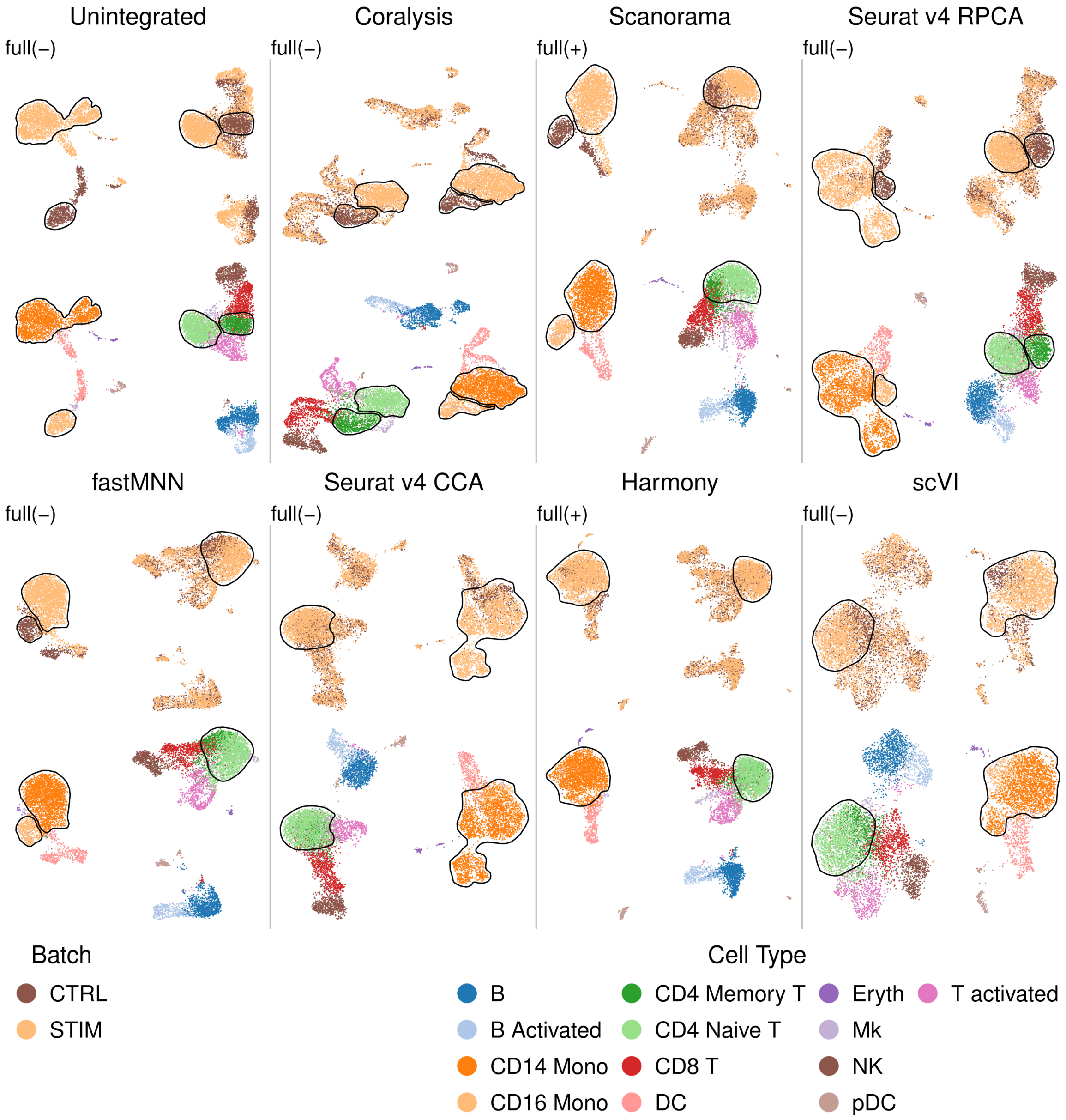


**Supplementary Figure 9 UMAP projections highlighting the batch and cell-type identities before and after integration of two PBMC scRNA-seq datasets through the scib-pipeline.** The two PBMC datasets consisted of one sample representing resting PBMCs (CTRL) and the other interferon-stimulated cells (STIM). CD4 naive T cells and CD14 monocytes were removed from the CTRL batch sample and CD4 memory T cells and CD16 monocytes from the STIM sample. Unshared similar cell type pairs are circumscribed by black lines. The best input-output combination was selected for every method. The label "full" represents input data with all features, and the minus and plus signs correspond to unscaled and scaled data, respectively.

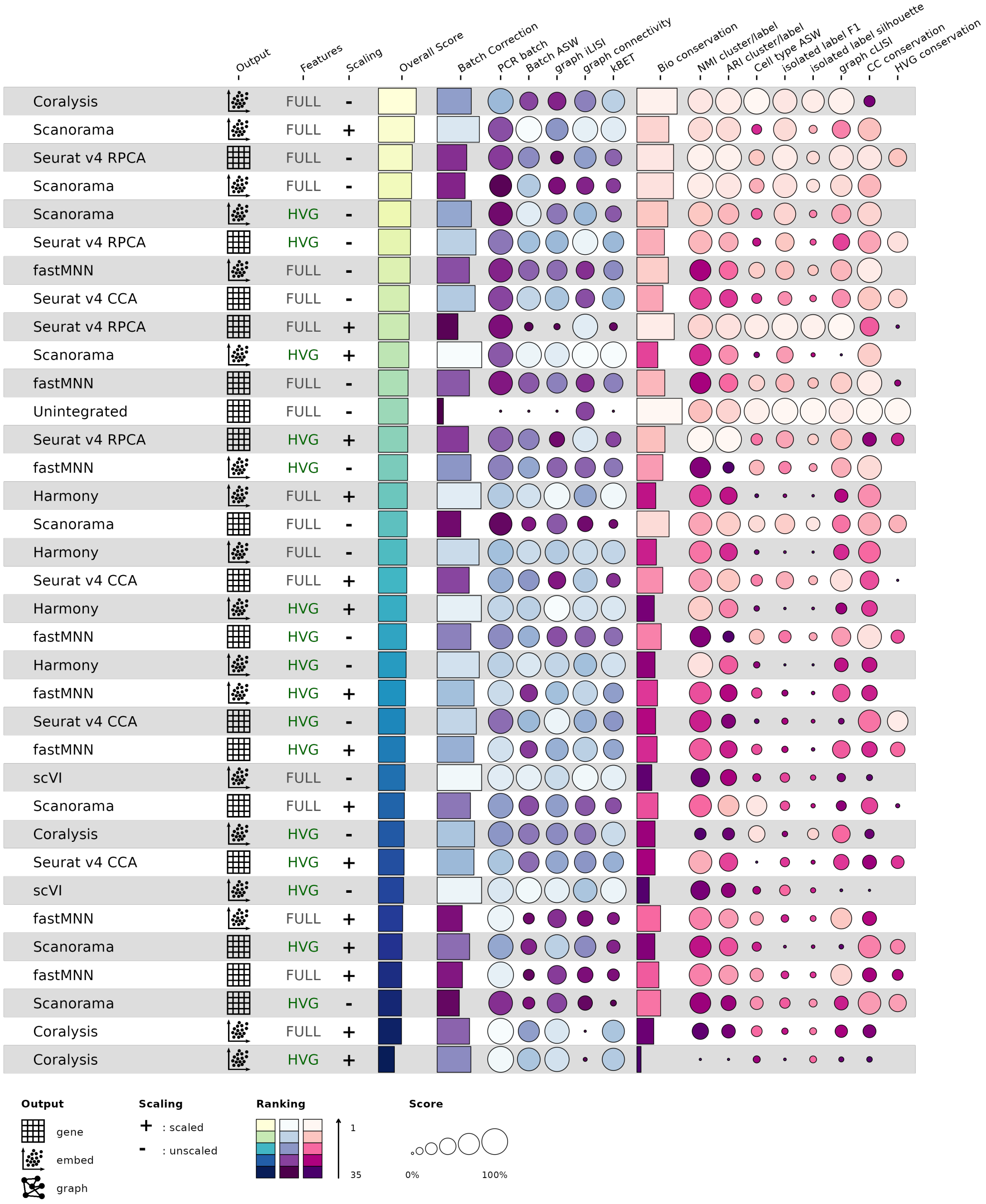


**Supplementary Figure 10 Performance ranking of integration methods by the overall score obtained with the scib-pipeline for the two PBMCs scRNA-seq data sets with unshared similar cell type pairs.** Overall score corresponds to 0.4:0.6 weighted mean between batch-correction (blue/purple) and bio-conservation (pink) metrics, respectively.

| Method | Bio conservation | | | | Batch correction | | | | | Aggregate score | | |
|---|---|---|---|---|---|---|---|---|---|---|---|---|
| | KMeans NMI | KMeans ARI | Silhouette label | cLISI | Silhouette batch | iLISI | KBET | Graph connectivity | PCR comparison | Batch correction | Bio conservation | Total |
| **Coralysis** | 0.76 | 0.60 | 0.66 | 1.00 | 0.89 | 0.24 | 0.45 | 0.82 | 0.94 | 0.67 | 0.75 | 0.72 |
| **Unintegrated** | 0.68 | 0.38 | 0.54 | 1.00 | 0.81 | 0.00 | 0.05 | 0.71 | 0.00 | 0.32 | 0.65 | 0.52 |

**Supplementary Figure 11 Assessment of integration performed with Coralysis on the ADT dataset provided by the scib-metrics python package using as ground-truth the cell-type labels given in Y. Hao et al. 2021 (at level 2 of granularity).**

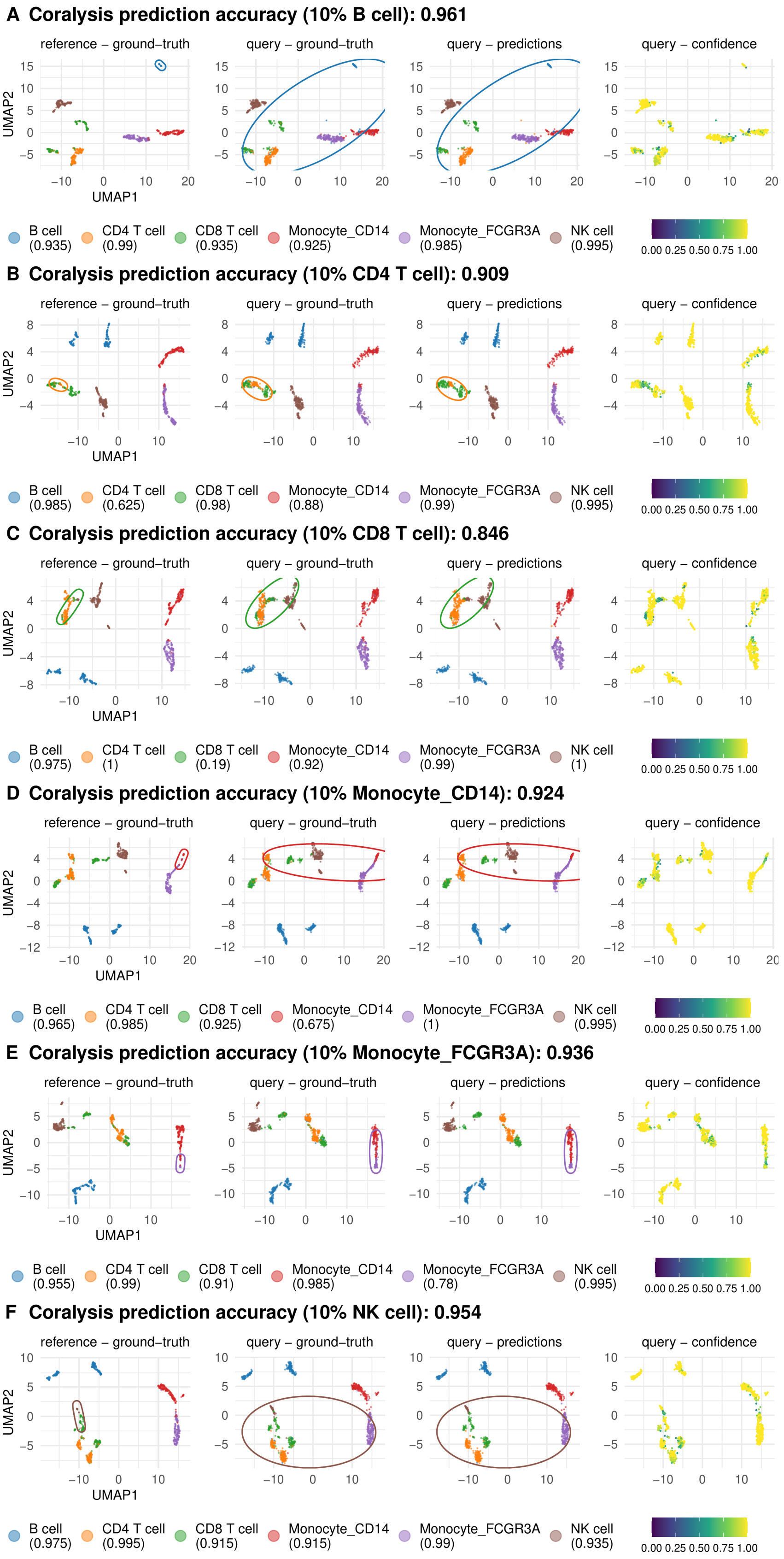
A Coralysis prediction accuracy (10% B cell): 0.961
reference – ground–truth
query – ground–truth
query – predictions
query – confidence
UMAP1
UMAP2
B cell (0.935)
CD4 T cell (0.99)
CD8 T cell (0.935)
Monocyte_CD14 (0.925)
Monocyte_FCGR3A (0.985)
NK cell (0.995)
0.00 0.25 0.50 0.75 1.00
B Coralysis prediction accuracy (10% CD4 T cell): 0.909
B cell (0.985)
CD4 T cell (0.625)
CD8 T cell (0.98)
Monocyte_CD14 (0.88)
Monocyte_FCGR3A (0.99)
NK cell (0.995)
C Coralysis prediction accuracy (10% CD8 T cell): 0.846
B cell (0.975)
CD4 T cell (1)
CD8 T cell (0.19)
Monocyte_CD14 (0.92)
Monocyte_FCGR3A (0.99)
NK cell (1)
D Coralysis prediction accuracy (10% Monocyte_CD14): 0.924
B cell (0.965)
CD4 T cell (0.985)
CD8 T cell (0.925)
Monocyte_CD14 (0.675)
Monocyte_FCGR3A (1)
NK cell (0.995)
E Coralysis prediction accuracy (10% Monocyte_FCGR3A): 0.936
B cell (0.955)
CD4 T cell (0.99)
CD8 T cell (0.91)
Monocyte_CD14 (0.985)
Monocyte_FCGR3A (0.78)
NK cell (0.995)
F Coralysis prediction accuracy (10% NK cell): 0.954
B cell (0.975)
CD4 T cell (0.995)
CD8 T cell (0.915)
Monocyte_CD14 (0.915)
Monocyte_FCGR3A (0.99)
NK cell (0.935)

**Supplementary Figure 12 Projection of query onto reference UMAP for every reference cell-type downsampled.** Projection of query onto reference UMAP highlighting ground-truth, predictions and confidence scores obtained with Coralysis reference-mapping method for every reference cell-type downsampled to 10%. The confidence scores represent the proportion of $K$ neighbors from the winning class ($K$=10). Ellipses circumscribe the position of the downsampled cell type.

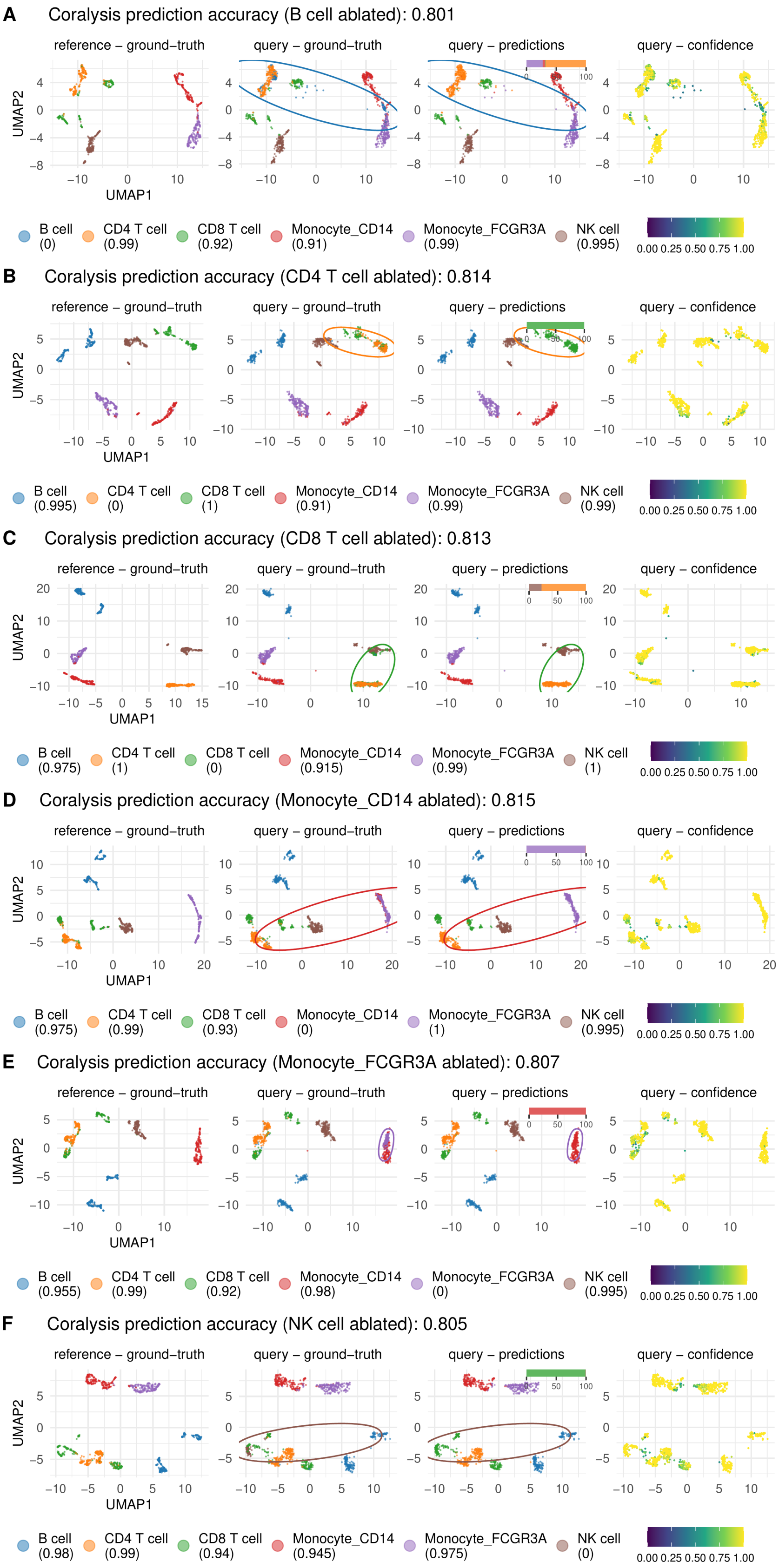
A
Coralysis prediction accuracy (B cell ablated): 0.801
reference – ground–truth
query – ground–truth
query – predictions
query – confidence
UMAP2
UMAP1
B cell (0)
CD4 T cell (0.99)
CD8 T cell (0.92)
Monocyte_CD14 (0.91)
Monocyte_FCGR3A (0.99)
NK cell (0.995)
0.00 0.25 0.50 0.75 1.00
B
Coralysis prediction accuracy (CD4 T cell ablated): 0.814
reference – ground–truth
query – ground–truth
query – predictions
query – confidence
UMAP2
UMAP1
B cell (0.995)
CD4 T cell (0)
CD8 T cell (1)
Monocyte_CD14 (0.91)
Monocyte_FCGR3A (0.99)
NK cell (0.99)
0.00 0.25 0.50 0.75 1.00
C
Coralysis prediction accuracy (CD8 T cell ablated): 0.813
reference – ground–truth
query – ground–truth
query – predictions
query – confidence
UMAP2
UMAP1
B cell (0.975)
CD4 T cell (1)
CD8 T cell (0)
Monocyte_CD14 (0.915)
Monocyte_FCGR3A (0.99)
NK cell (1)
0.00 0.25 0.50 0.75 1.00
D
Coralysis prediction accuracy (Monocyte_CD14 ablated): 0.815
reference – ground–truth
query – ground–truth
query – predictions
query – confidence
UMAP2
UMAP1
B cell (0.975)
CD4 T cell (0.99)
CD8 T cell (0.93)
Monocyte_CD14 (0)
Monocyte_FCGR3A (1)
NK cell (0.995)
0.00 0.25 0.50 0.75 1.00
E
Coralysis prediction accuracy (Monocyte_FCGR3A ablated): 0.807
reference – ground–truth
query – ground–truth
query – predictions
query – confidence
UMAP2
UMAP1
B cell (0.955)
CD4 T cell (0.99)
CD8 T cell (0.92)
Monocyte_CD14 (0.98)
Monocyte_FCGR3A (0)
NK cell (0.995)
0.00 0.25 0.50 0.75 1.00
F
Coralysis prediction accuracy (NK cell ablated): 0.805
reference – ground–truth
query – ground–truth
query – predictions
query – confidence
UMAP2
UMAP1
B cell (0.98)
CD4 T cell (0.99)
CD8 T cell (0.94)
Monocyte_CD14 (0.945)
Monocyte_FCGR3A (0.975)
NK cell (0)
0.00 0.25 0.50 0.75 1.00

**Supplementary Figure 13 Projection of query onto reference UMAP for every reference cell-type ablated.** Projection of query onto reference UMAP highlighting ground-truth, predictions and confidence scores obtained with Coralysis reference-mapping method for every reference cell-type downsampled to 10%. The bar plot above the UMAP corresponds to the cell-type labels against which the query cell-type, ablated in the reference, was classified. The confidence scores represent the proportion of $K$ neighbors from the winning class ($K$=10). Ellipses circumscribe the position of the ablated cell type.

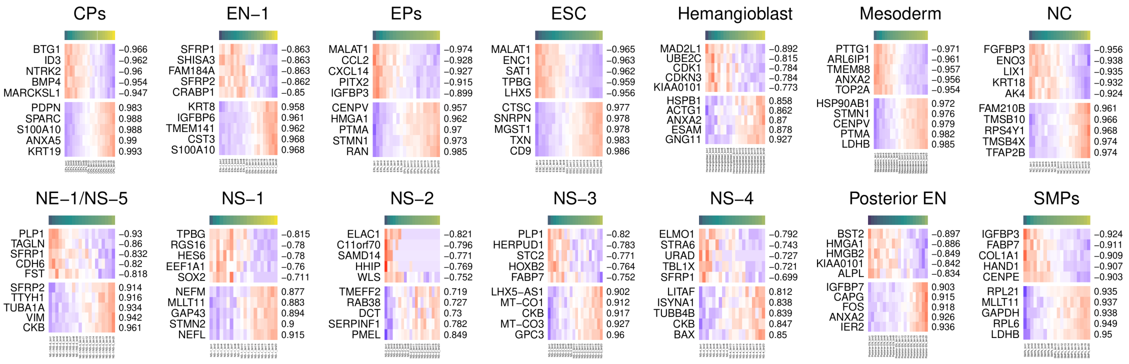


**Supplementary Figure 14 Expression of the top five negative and positive correlated genes across the Coralysis cell cluster probability bins for each embryoid body cell-type.** Pearson correlation was performed using the mean cell cluster probability and the average gene expression across the 20 bins of Coralysis cell cluster probability. Top color bar corresponds to the mean cell cluster probability per bin. The averaged gene expression was represented by Z-scores (by row).
